# ORP9-PH domain-based fluorescent reporters for visualizing phosphatidylinositol 4-phosphate dynamics in living cells

**DOI:** 10.1101/2023.12.14.571782

**Authors:** Moeka Ajiki, Masaru Yoshikawa, Tomoki Miyazaki, Asami Kawasaki, Kazuhiro Aoki, Fubito Nakatsu, Shinya Tsukiji

## Abstract

Fluorescent reporters that visualize phosphatidylinositol 4-phosphate (PI4P) in living cells are indispensable to elucidate the roles of this fundamental lipid in cell physiology. However, currently available PI4P reporters have limitations, such as Golgi-biased localization and low detection sensitivity. Here, we present a series of fluorescent PI4P reporters based on the pleckstrin homology (PH) domain of oxysterol-binding protein-related protein 9 (ORP9). We show that the green fluorescent protein AcGFP1-tagged ORP9-PH domain can be used as a fluorescent PI4P reporter to detect cellular PI4P across its wide distribution at multiple cellular locations, including the plasma membrane (PM), Golgi, endosomes, and lysosomes with high specificity and contrast. We also developed blue, red, and near-infrared fluorescent PI4P reporters suitable for multicolor fluorescence imaging experiments. Finally, we demonstrate the utility of the ORP9-PH domain-based reporter to visualize dynamic changes in the PI4P distribution and level in living cells upon synthetic ER–PM membrane contact manipulation and GPCR stimulation. This work offers a new set of genetically encoded fluorescent PI4P reporters that are practically useful for the study of PI4P biology.

## Introduction

Phosphoinositides are a class of low-abundance phospholipids in cellular membranes that play key regulatory roles in cell physiology.^1–3^ Their inositol headgroup is reversibly phosphorylated or dephosphorylated at the 3, 4, or 5 position of the inositol ring by phosphoinositide kinases or phosphatases.^4^ Phosphatidylinositol 4-phosphate (PI4P) is one of the most abundant species of phosphoinositides in mammalian cells.^1,5–7^ The subcellular distribution of PI4P is spatially controlled by its metabolic enzymes, which are localized at distinct cellular membranes. PI4P is synthesized *de novo* by phosphatidylinositol 4-kinases (PI4Ks) at the plasma membrane (PM), Golgi, endosomes, and lysosomes.^8,9^ A pool of PI4P synthesized at these membranes is transported to the ER through membrane contact sites where the PM or organelle membranes are closely apposed.^10^ At these membrane contact sites, oxysterol-binding protein (OSBP)-related proteins (ORPs),^11^ a family of so-called lipid transfer proteins,^12,13^ mediate the transfer of PI4P to the ER. The transferred PI4P is then metabolically degraded by Sac1, an ER-resident PI4P phosphatase, at the ER.^3,14–18^ This spatially controlled metabolic pathway mainly determines the distribution and abundance of PI4P within the cells. From a functional standpoint, the distribution and levels of PI4P are critical to its fundamental roles in cell physiology, which include but are not limited to roles in lipid metabolism, membrane trafficking, and signaling.^1–3,5,8^ Moreover, recent studies have revealed roles of PI4P in such functions as membrane contact site organization,^14,15,19-22^ autophagy,^23-28^ lysosome repair,^29^ pathogen lifecycle,^30,31^ and inflammatory signaling.^32,33^

To elucidate the role of PI4P in regulating cellular physiology, fluorescent reporters that visualize the dynamic change of PI4P distribution and level in living cells are indispensable.^5,34,35^ For this purpose, fluorescent protein fusions of PI4P-binding pleckstrin homology (PH) domains derived from OSBP and FAPP1 have been developed and widely used.^36,37^ However, these probes show biased detection of Golgi PI4P due to additional interactions with ADP-ribosylation factor 1 (Arf1), a Golgi-resident protein,^36,38^ and are not suited for detecting global PI4P localization or the PM pools of this lipid. To overcome these issues, a unique PI4P-binding domain of the SidM protein (P4M) derived from the pathogen *Legionella pneumophila* was used to construct fluorescent PI4P reporters.^39^ P4M-based reporters exhibit an improved ability to detect PI4P across its wide distribution at multiple cellular locations, including the Golgi, PM, and late endosomes/lysosomes. However, the reporters show low-to-moderate detection sensitivity depending on cell types, leading to high background (cytosolic) signals. Therefore, the development of fluorescent reporters for unbiased and sensitive detection of PI4P dynamics in living cells is still in high demand.

Here we present a series of new fluorescent PI4P reporters based on the PH domain of ORP9. ORP9 is a member of the ORP family and mediates the transfer of cholesterol or exchange of PI4P and phosphatidylserine (PS) at membrane contact sites.^15,40^ ORP9 contains a PI4P-specific PH domain,^40^ but thus far the ORP9-PH domain has not been used for constructing PI4P reporters. In this study, we show that a green fluorescent protein fusion of the ORP9-PH domain is capable of visualizing cellular PI4P across its distribution at multiple cellular locations including the PM, Golgi, endosomes, and lysosomes with high specificity and contrast. We then show that the use of fluorescent proteins with a (weak) dimerization tendency is critical to creating sensitive PI4P reporters. Using this information as a guideline, we successfully generate an assortment of blue, green, red, and near-infrared fluorescent PI4P reporters suitable for multicolor fluorescence imaging experiments. We further demonstrate that the ORP9-PH-based reporter can be used to monitor dynamic changes in the PI4P level in living cells upon synthetic ER–PM membrane contact manipulation and GPCR stimulation.

## Results and discussion

### Design of novel fluorescent PI4P reporters

To develop novel fluorescent PI4P reporters, we focused on the PH domain of ORP9. ORP9 is localized at membrane contact sites between the ER and the Golgi or endo-lysosomes, where it mediates the transfer of lipids.^15,40^ It was previously reported that ORP9 possesses a PI4P-specific PH domain at its N-terminus.^40^ In the course of our previous study, we realized that the ORP9-PH domain can be a useful PI4P reporter.^41^ We therefore attempted to determine whether the ORP9-PH domain can be used to generate a novel fluorescent PI4P reporter with unbiased and sensitive detection properties. To this end, we first evaluated the binding affinity of the recombinant ORP9-PH domain to PI4P *in vitro*. Using isothermal titration calorimetry (ITC), we determined the dissociation constant (*K*_D_) to be 0.94 µM (**Figure S1**), which was slightly larger than that of the OSBP-PH domain (250 nM) but was almost comparable to that of the P4M domain (1 µM).^34^

### The AcGFP1-tagged ORP9-PH domain visualizes PI4P in living cells

Next, we attempted to construct a fluorescent PI4P reporter based on the ORP9-PH domain. For this purpose, we chose AcGFP1 as a green fluorescent protein and fused it to the N-terminus of the ORP9-PH domain (AcGFP1-ORP9-PH). We then transfected HeLa cells with AcGFP1-ORP9-PH. Confocal fluorescence imaging showed that AcGFP1-ORP9-PH was distributed to the PM and other organelles (**Figure 1a**). To identify the organelles to which AcGFP1-ORP9-PH localized, we performed colocalization assays using organelle markers. The results revealed that in addition to the PM, AcGFP1-ORP9-PH was localized to the Golgi, lysosomes, and Rab7-positive late endosomes (**Figure 1a**). AcGFP1-ORP9-PH also colocalized with a subset of Rab5-positive early endosomes (**Figure 1a**). All these results are consistent with the previously reported intracellular distribution of PI4P.^3,5,8,39,41^

**Figure 1.**
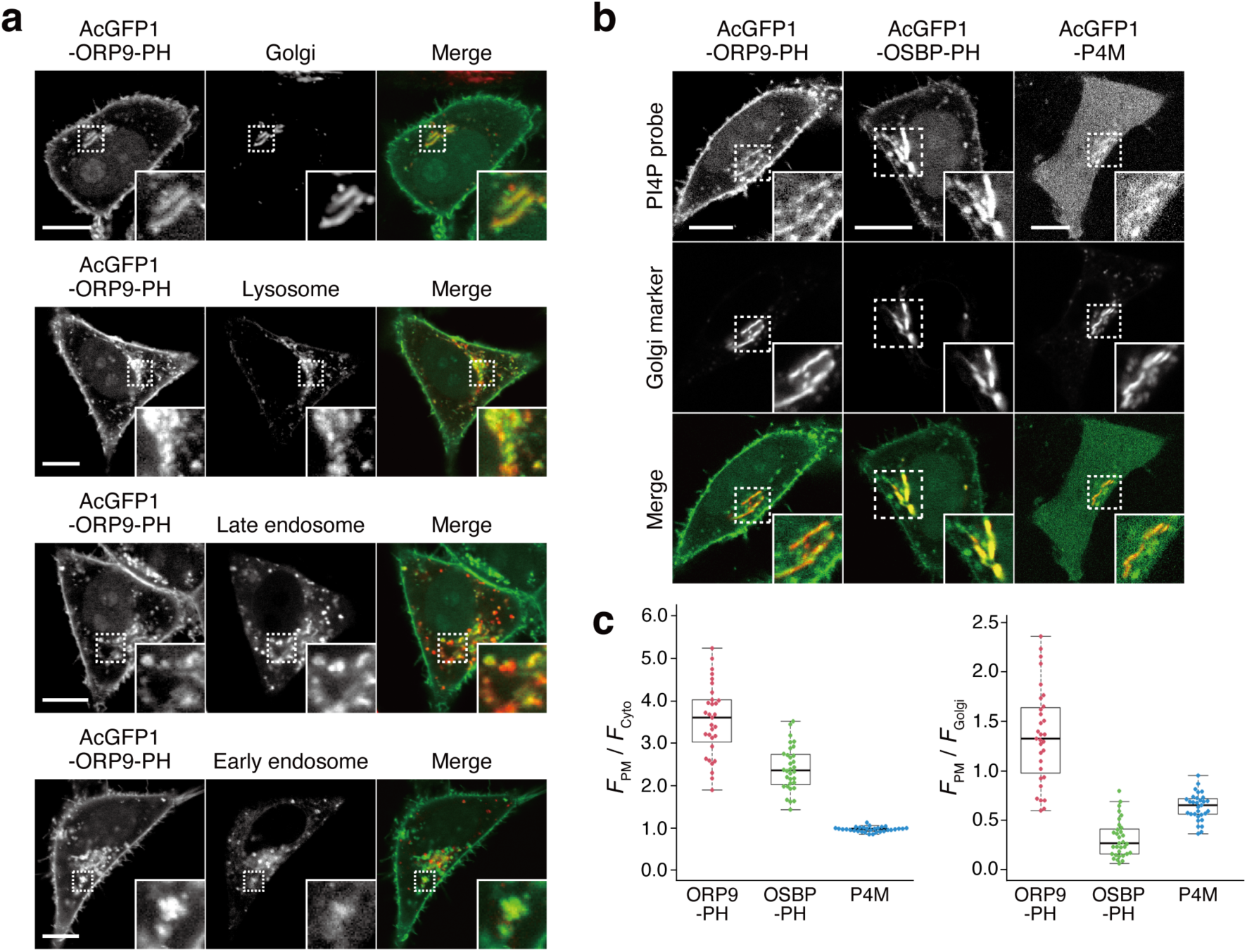
Intracellular distribution of AcGFP1-ORP9-PH and other PI4P probes. (**a**) Confocal fluorescence images of HeLa cells coexpressing AcGFP1-ORP9-PH and organelle markers: the Golgi apparatus, GalT-mCherry; lysosome, LAMP1-mCherry; late endosome, iRFP713-Rab7; early endosome: iRFP713-Rab5. Insets show the regions indicated by white dashed boxes at higher magnification. Scale bars, 10 μm. (**b**) Comparison of AcGFP1-ORP9-PH, AcGFP1-OSBP-PH, and AcGFP1-P4M. Confocal fluorescence images of HeLa cells coexpressing each PI4P probe and Golgi marker (GalT-mCherry) are shown. Insets show the regions indicated by white dashed boxes at higher magnification. Scale bars, 10 μm. Additional images are shown in **Figure S3**. (**c**) Quantification of the PI4P detection sensitivity and Golgi bias. The PI4P detection sensitivity was evaluated by quantifying the ratios of the PM to the cytosolic fluorescence intensity of PI4P reporters (left). The Golgi bias was evaluated by quantifying the ratios of the PM to the Golgi fluorescence intensity of PI4P reporters (right). Central lines represent the median values (n ≥ 30 cells).

The PH domains of some ORP family members, such as ORP5 and ORP8, have been reported to bind to PI(4,5)P_2_ as well as PI4P.^21,22,42^ We thus performed lipid depletion experiments to evaluate the PI4P specificity of the ORP9-PH domain-based reporter. We first used A1, a small-molecule inhibitor of PI4KIIIα, to deplete the PM pools of PI4P specifically.^21,43,44^ Upon treatment of HeLa cells expressing AcGFP1-ORP9-PH with A1 (100 nM) for 20 min, AcGFP1-ORP9-PH was completely dissociated from the PM, whereas a PI(4,5)P_2_ reporter (mTagBFP2-PLCδ-PH)^44^ showed almost no change in PM concentration (**Figure S2a-d**). To further confirm that the ORP9-PH domain-based reporter does not bind to PI(4,5)P_2_, we depleted PM PI(4,5)P_2_ by using a rapamycin inducible INPP5E [PI(4,5)P_2_ phosphatase] recruitment system (**Figure S2e**).^44^ Despite significant depletion of the PM pools of PI(4,5)P_2_, which was monitored by the PI(4,5)P_2_ reporter, no dissociation of AcGFP1-ORP9-PH was observed (**Figure S2f**). Rather, the PM concentration of AcGFP1-ORP9-PH was slightly but significantly increased by PM PI(4,5)P_2_ depletion, as indicated by the PM dissociation index of 0.90 (a PM dissociation index less than 1.0 indicates PM association of AcGFP1-ORP9-PH) (**Figure S2h**). This result can be ascribed to the increase in the PM PI4P level due to the INPP5E-mediated dephosphorylation of PM PI(4,5)P_2_ at the 5’ position, which results in further recruitment of AcGFP1-ORP9-PH to the PM. Overall, these colocalization and lipid depletion assays demonstrate that AcGFP1-ORP9-PH can be used as a fluorescent reporter for specifically detecting cellular PI4P across its wide distribution at the PM and other organelles, including the Golgi, endosomes, and lysosomes.

### Comparison of the ORP9-PH domain-based reporter with previous PI4P reporters

To further characterize the ORP9-PH domain-based PI4P reporter, we experimentally compared the live-cell PI4P-detection properties of AcGFP1-ORP9-PH and previous OSBP-PH domain- and P4M domain-based PI4P reporters. We used the same AcGFP1 as a fluorescent protein for OSBP-PH domain- (AcGFP1-OSBP-PH) and P4M domain-based (AcGFP1-P4M) reporters to make a fair comparison of the properties of each PI4P-binding domain. In transient expression experiments using HeLa cells, AcGFP1-ORP9-PH and AcGFP1-OSBP-PH detected cellular PI4P with lower background (cytosolic) signals than AcGFP1-P4M (**Figure 1b,c** and **Figure S3a**). Moreover, although AcGFP1-OSBP-PH showed a sufficient PI4P-detection ability, the reporter was strongly biased to localize to the Golgi membrane, as previously reported (**Figure 1b,c** and **Figure S3b**).^30,31^ In contrast, AcGFP1-ORP9-PH did not exhibit such Golgi-biased localization and enabled the detection of PM PI4P more efficiently than AcGFP1-OSBP-PH and AcGFP1-P4M (**Figure 1b,c** and **Figure S3c**). The same trend was also observed in other cell lines such as Cos-7 and HEK293 cells (**Figure S4**).^45^ Taken together, these results indicate the utility of AcGFP1-ORP9-PH as a novel, practical, bias-less fluorescent reporter for visualizing the cellular distribution of PI4P with high sensitivity and contrast.

### Expansion of the color palette of ORP9-PH domain-based PI4P reporters

Having successfully developed the green fluorescent PI4P reporter, AcGFP1-ORP9-PH, we next moved on to generate its color variants. We first attempted to construct a red fluorescent PI4P reporter by using mScarlet-I^46^ as a red fluorescent protein. We fused mScarlet-I to the N-terminus of the ORP9-PH domain (mScarlet-I-ORP9-PH) and expressed the protein in HeLa cells. Unexpectedly, mScarlet-I-ORP9-PH showed no or only low PI4P detection ability (**Figure 2b,c** and **Figure S5a**). However, when we used dTomato^47^ as an alternative red fluorescent protein (dTomato-ORP9-PH), dTomato-ORP9-PH was distributed to the PM, Golgi, lysosomes, and endosomes with high contrast, as with the case of AcGFP1-ORP9-PH (**Figure 2a,c** and **Figure S6**). The PI4P-dependent membrane localization of dTomato-ORP9-PH was also confirmed by lipid depletion experiments using A1 (**Figure S6c**), demonstrating that dTomato-ORP9-PH serves as a red fluorescent PI4P reporter.

**Figure 2.**
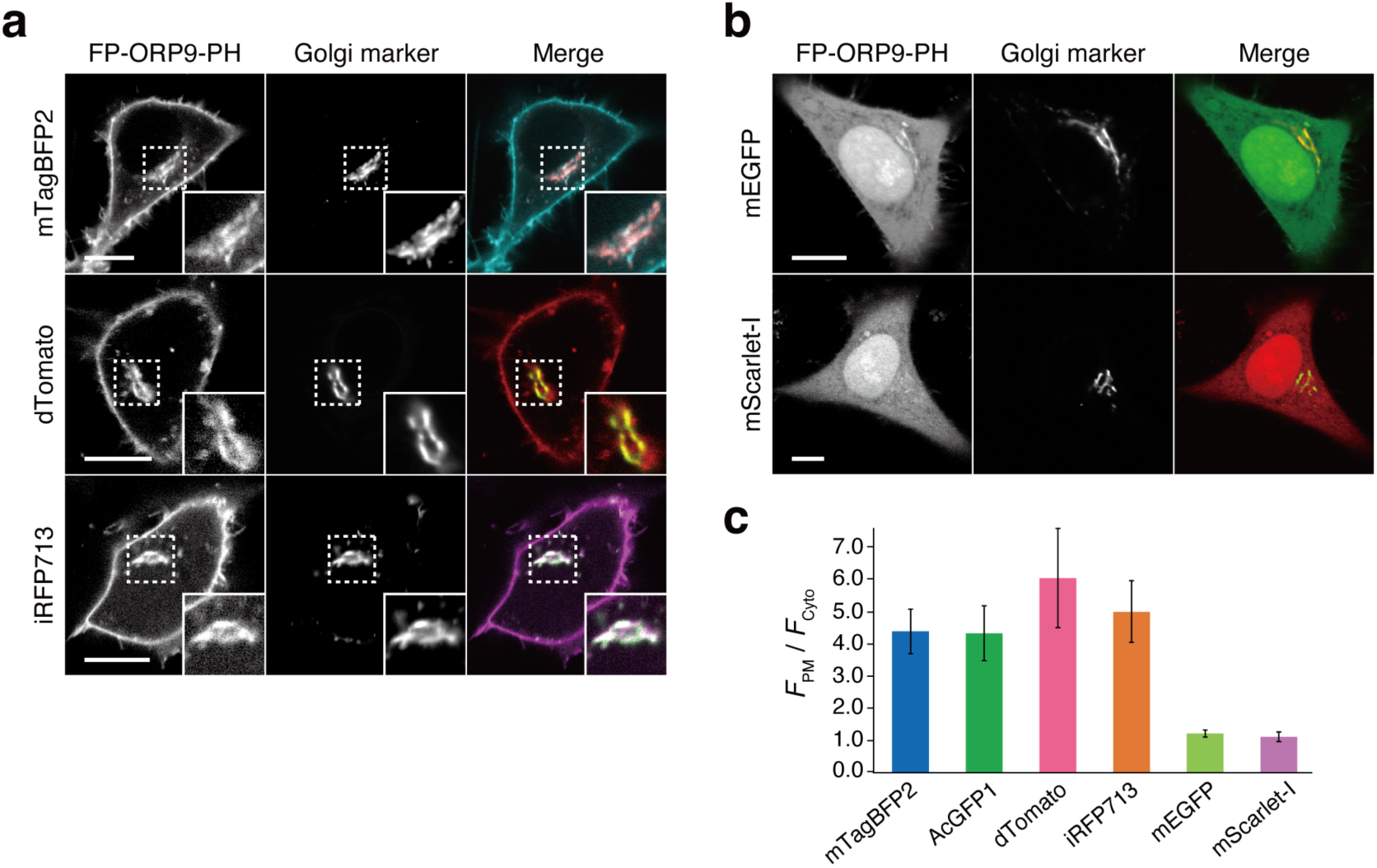
Expansion of the color palette of ORP9-PH domain-based PI4P probes. (**a**) Confocal fluorescence images of HeLa cells coexpressing dimeric fluorescent protein (FP)-tagged ORP9-PH domains [mTagBFP2-ORP9-PH (top), dTomato-ORP9-PH (middle), and iRFP713-ORP9-PH (bottom)] and the Golgi marker (GalT-mCherry for mTagBFP2-ORP9-PH and iRFP713-ORP9-PH; GalT-AcGFP1 for dTomato-ORP9-PH). Insets show at higher magnification the regions indicated by white dashed boxes. Scale bars, 10 μm. Additional images are shown in **Figures S6, S8 and S9**. (**b**) Confocal fluorescence images of HeLa cells coexpressing monomeric FP-tagged ORP9-PH domains [mEGFP-ORP9-PH (top) and mScarlet-I-ORP9-PH (bottom)] and the Golgi marker (GalT-mCherry for mEGFP-ORP9-PH; GalT-AcGFP1 for mScarlet-I-ORP9-PH). Scale bars, 10 μm. Additional images are shown in **Figure S5**. (**c**) Quantification of the PI4P detection sensitivity. The PI4P detection sensitivity was evaluated by quantifying the ratios of the PM to the cytosolic fluorescence intensity (*F*_PM_/*F*_cyto_) of FP-tagged ORP9-PH domains: mTagBFP2-ORP9-PH (blue), AcGFP1-ORP9-PH (green), dTomato-ORP9-PH (pink), iRFP713-ORP9-PH (orange), mEGFP-ORP9-PH (light green), and mScarlet-I-ORP9-PH (purple). Data are presented as the mean ± SD (n = 15 cells).

The contrasting results for the mScarlet-I and dTomato fusion proteins described above imply that the choice of fluorescent proteins is critical to engineering practically useful ORP9-PH domain-based PI4P reporters. As a key factor determining this issue, we focused on the dimerization tendency of fluorescent proteins. While mScarlet-I is monomeric,^46^ dTomato is dimeric.^47^ Moreover, according to the guideline of the OSER (organized smooth endoplasmic reticulum) assay,^48–50^ the green fluorescent protein AcGFP1, which has a reported OSER score of 29.5%,^49^ is likely to be dimeric. Based on these observations, we hypothesized that dimeric fluorescent proteins are more suited as fluorescent tags for the construction of ORP9-PH domain-based PI4P reporters than monomeric fluorescent proteins. To investigate this hypothesis, we replaced AcGFP1 of AcGFP1-ORP9-PH with a monomeric green fluorescent protein mEGFP^51^ (mEGFP-ORP9-PH). As in the case of mScarlet-I-ORP9-PH, mEGFP-ORP9-PH exhibited no or very low levels of PI4P detection ability (**Figure 2b,c** and **Figure S5b**), which was consistent with our hypothesis. We further tested a tetrameric fluorescent protein, AzamiGreen,^52^ as a fluorescent tag (AzamiGreen-ORP9-PH). In this case, AzamiGreen-ORP9-PH formed unexpected fluorescent clusters in the cell and showed almost no localization to the Golgi (**Figure S7a**). In addition, AzamiGreen-ORP9-PH did not dissociate completely from the PM even after the cells were treated with A1 (**Figure S7b**), indicating its unsuitability as a PI4P reporter. These observations can probably be ascribed to the avidity effect. That is, the PI4P-binding affinity of the monomeric fluorescent protein fusion is too weak to detect cellular PI4P with sufficient efficiency, while that of the tetrameric fluorescent protein fusion is too high, causing inhibition of PI4P metabolism. Based on the collective results of these experiments, the use of dimeric fluorescent proteins, but neither monomeric nor tetrameric fluorescent proteins, is strongly recommended for constructing ORP9-PH domain-based PI4P reporters.

Following the above design principles, we further engineered blue and near-infrared fluorescent PI4P reporters by using mTagBFP2^53^ (mTagBFP2-ORP9-PH), which can be classified as a weak dimer based on its reported OSER score of 49.8%,^49,50,54^ and the dimeric iRFP713^55^ (iRFP713-ORP9-PH), respectively. Colocalization assays and lipid depletion experiments verified that both mTagBFP2-ORP9-PH and iRFP713-ORP9-PH function as reliable reporters to visualize the PI4P distribution in cells (**Figure S8,9**).

### Visualization of PI4P dynamics in living cells

Finally, we investigated whether the ORP9-PH domain-based fluorescent reporter can be used to visualize the spatiotemporal dynamics of PI4P in living cells. We first took a synthetic biology approach in which the PM PI4P level was artificially modulated by chemically controlling lipid transport at ER–PM contact sites. It was previously demonstrated that acute tethering of ER-bound ORP5 to the PM using a rapamycin-induced protein dimerization tool triggers PM PI4P depletion by the PI4P/phosphatidylserine (PS) exchange activity of ORP5.^21^ Accordingly, we constructed a synthetic system that allows the control of the PM tethering of ORP5 (and thus PM PI4P depletion) in a reversible manner using our previously developed small-molecule trimethoprim (TMP)-HaloTag ligand conjugate (TMP-HL) (**Figure S10**), which serves as a chemical dimerizer for *Escherichia coli* dihydrofolate reductase (eDHFR) and HaloTag.^56,57^ In this system, an engineered ORP5 in which the N-terminal PH domain was replaced with mScarlet-I-tagged eDHFR [mScarlet-I-eDHFR-ORP5(ΔPH)] was expressed at the ER, while HaloTag fused to the C-terminal PM-targeting motif of KRas4B [HaloTag-KRas4B(CT)] was expressed at the inner leaflet of the PM (**Figure 3a)**. The PM tethering of the engineered ORP5 can be induced by the addition of TMP-HL, and can also be reversed by the subsequent addition of (unmodified) TMP as a competitor ligand. To monitor PI4P dynamics controlled by synthetic ORP5-mediated ER–PM contact formation, we co-expressed mScarlet-I-eDHFR-ORP5(ΔPH), HaloTag-KRas4B(CT), and AcGFP1-ORP9-PH in HeLa cells. At the initial state, mScarlet-I-eDHFR-ORP5(ΔPH) was widely distributed to the ER membrane and AcGFP1-ORP9-PH was localized at the PM (**Figure 3b**). Upon TMP-HL addition, mScarlet-I fluorescence was observed as discontinuous puncta associated with the PM, demonstrating the formation of ER–PM contact sites via PM tethering of mScarlet-I-eDHFR-ORP5(ΔPH). Along with this ER–PM contact formation, AcGFP1-ORP9-PH was dissociated from the PM over 20 min [with a half-maximal translocation time (*t*_1/2_) of 7 min] and the PM PI4P-depleted state was stably maintained at least for 80 min (**Figure 3c,e**). However, treating cells with TMP after TMP-HL addition led to the disassembly of ER–PM contacts and relocalization of AcGFP1-ORP9-PH to the PM over 1 h (with a *t*_1/2_ of 13 min) (**Figure 3b,e**). Control experiments using DMSO showed no change in the localization of mScarlet-I-eDHFR-ORP5(ΔPH) and AcGFP1-ORP9-PH, as expected (**Figure 3d,e**). These observations demonstrate the ability of the ORP9-PH domain-based fluorescent reporter to visualize PM PI4P depletion and replenishment induced by synthetic control of ORP5-mediated ER–PM contact formation.

**Figure 3.**
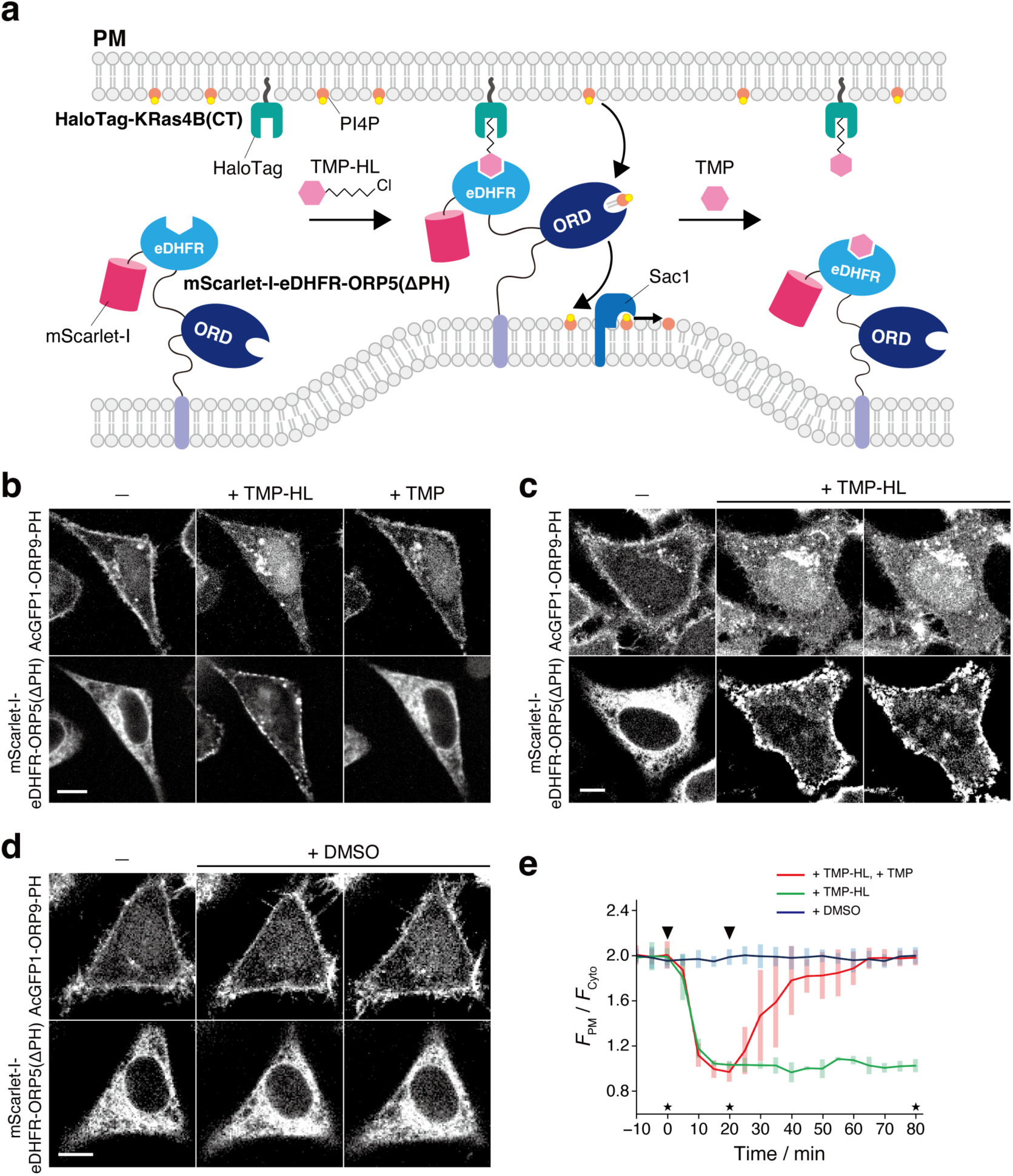
Visualization of PI4P dynamics with AcGFP1-ORP9-PH upon synthetic ORP5-mediated ER–PM contact manipulation. (**a**) Schematic illustration of the experimental setup. (**b**) Confocal fluorescence images of HeLa cells coexpressing mScarlet-I-eDHFR-ORP5(ΔPH), HaloTag-KRas4B(CT), and AcGFP1-ORP9-PH. Images were taken at the time points indicated by the asterisks in panel **e**: before (left) and 20 min after the addition of TMP-HL (0.5 µM) (center), and 60 min after the subsequent addition of TMP (100 µM). Scale bar, 10 μm. For the time-lapse movie, see **Movie S1**. (**c**,**d**) Control experiments for **b**. The experiments were performed in the same manner as in **b** except that cells were treated with TMP-HL only (**c**) or DMSO (**d**). Scale bars, 10 μm. (**e**) Time course of the PM PI4P level monitored by AcGFP1-ORP9-PH. The ratios of the PM to the cytosolic fluorescence intensity (*F*_PM_/*F*_cyto_) of AcGFP1-ORP9-PH are plotted as a function of time. The arrowheads in the graph indicate the time of TMP-HL (or DMSO) and TMP (or none) addition. Data are presented as the mean ± SD (n = 5 cells).

In the field of phosphoinositide signaling, there has been a growing interest in the role of PI4P in GPCR signaling.^5,58,59^ Recent studies revealed that a PM pool of PI4P can be depleted upon GPCR stimulation.^60^ Next, therefore, we attempted to apply the ORP9-PH domain-based fluorescent reporter to monitor the dynamic change of PI4P following GPCR activation. Here we targeted the muscarinic M1 receptor (M1R) and co-expressed this protein and AcGFP1-ORP9-PH in HeLa cells. Upon stimulation with the M1R agonist carbachol,^60^ AcGFP1-ORP9-PH, which was observed at the PM at the initial state, was immediately dissociated from the PM with a *t*_1/2_ of <1 min (**Figure 4a,d**). Furthermore, when the M1R antagonist atropine was subsequently added, AcGFP1-ORP9-PH returned to the PM with a t_1/2_ of 2 min (**Figure 4a,d**). No relocalization of AcGFP1-ORP9-PH to the PM was observed for at least 20 min when the cells were treated only with carbachol, i.e., without subsequent atropine (**Figure 4b,d**). DMSO treatment induced no change in the PM PI4P level (**Figure 4c,d**). These results showcase that the ORP9-PH domain-based fluorescent reporter is practically useful for visualizing the rapid dynamic change of the PM PI4P level upon GPCR stimulation.

**Figure 4.**
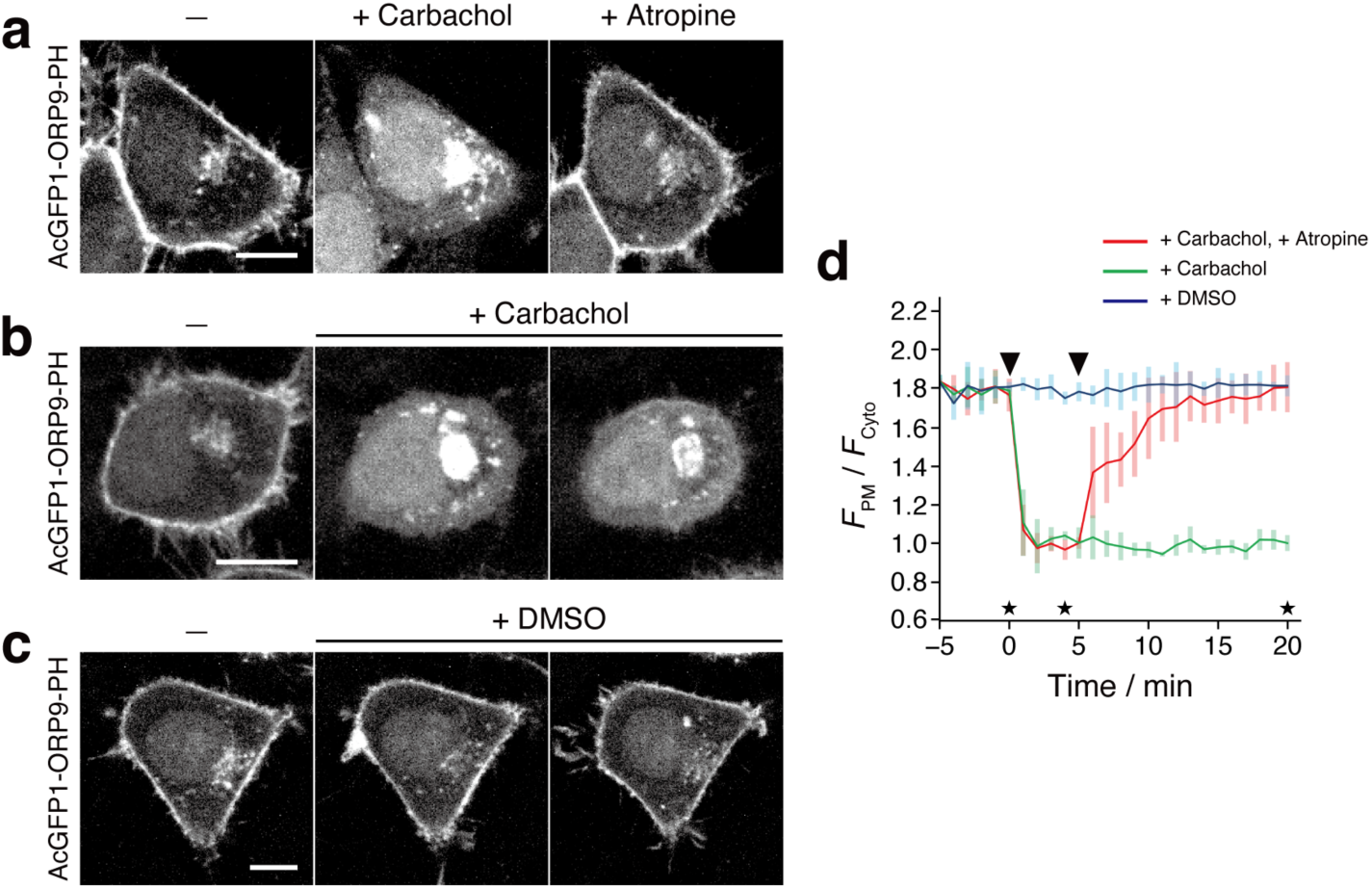
Visualization of PI4P dynamics with AcGFP1-ORP9-PH upon M1R stimulation. (**a**) Confocal fluorescence images of HeLa cells coexpressing M1R and AcGFP1-ORP9-PH. Images were taken at the time points indicated by the asterisks in panel **d**: before (left) and 4 min after the addition of carbachol (1 mM) (center), and 15 min after the subsequent addition of atropine (10 µM). Scale bar, 10 μm. For the time-lapse movie, see **Movie S2**. (**b**,**c**) Control experiments for **a**. The experiments were performed in the same manner as in **a** except that cells were treated with carbachol only (**b**) or DMSO (**c**). Scale bars, 10 μm. (**d**) Time course of the PM PI4P level monitored by AcGFP1-ORP9-PH. The ratios of the PM to the cytosolic fluorescence intensity (*F*_PM_/*F*_cyto_) of AcGFP1-ORP9-PH are plotted as a function of time. The arrowheads in the graph indicate the time of carbachol (or DMSO) and atropine (or none) addition. Data are presented as the mean ± SD (n ≥ 5 cells).

### Conclusions

In this work, we demonstrated that the ORP9-PH domain can be used as a novel PI4P-specific binding motif to construct fluorescent PI4P reporters. Using fluorescent proteins with a (weak) dimerization tendency, we developed a set of blue, green, red, and near-infrared fluorescent PI4P reporters, all of which were capable of specifically visualizing cellular PI4P distributed at the PM, Golgi, endosome, and lysosome membranes with high sensitivity and contrast. It is also noteworthy that ORP9-PH domain-based reporters are particularly suitable for detecting the PM pool of PI4P, which has often proven difficult using the previously developed PI4P reporters. We further demonstrated the applicability of the green fluorescent ORP9-PH domain-based reporter for visualizing the dynamic change of PI4P in living cells upon synthetic ER–PM contact manipulation and agonist-induced GPCR activation.

The present fluorescent protein-tagged ORP9-PH domains have a limitation common to other translocation-based reporters. As observed in several experiments, PM-localized PI4P reporters relocalized not only to the cytosol but also to other PI4P-containing organelle membranes, including that of the Golgi, when PM PI4P was acutely depleted. This is because the present reporters are in equilibrium with each organelle membrane-bound pool and an unbound cytosolic pool, and a change in the PI4P level at one location can affect the overall reporter localization.^5^ Therefore, care needs to be taken when analyzing the dynamic change of the PI4P level at specific locations upon cell stimulation. This issue may be overcome by converting the present single fluorophore-, translocation-based reporters to membrane-targeted FRET-based reporters,^61–63^ which is currently underway.

In conclusion, the ORP9-PH-based fluorescent PI4P reporters developed in this work will serve as useful new tools for studying the dynamics and functions of PI4P in regulating cell physiology.

## Materials and methods

### Plasmid construction

All the cDNA and amino acid sequences of the constructs used in this study are listed in the “**Supplementary Sequences**” section. pEGFP-N1 (Clontech), pEGFP-C1 (Clontech), and pCAGGS^64^ (provided by Dr. Jun-ichi Miyazaki, Osaka University) were used as vector backbones. pPBbsr^65^ (blasticidin S resistance) was used as a piggyBac donor vector for the establishment of stable cell lines. The following plasmids were purchased or kindly provided from the following companies or researchers, and used directly or as PCR templates to construct expression plasmids: mCherry-ORP9,^41^ GFP-OSBP-PH,^41^ and mRFP-FKBP-ΔPH-ORP5 ^21^ from F.N.; pAcGFP1-Golgi from Clontech; mCherry-P4M-SidM (Addgene plasmid #51471),^39^ iRFP-FRB-Rab5 (Addgene plasmid #51612),^39^ and iRFP-FRB-Rab7 (Addgene plasmid #51613)^39^ from Dr. Tamas Balla (National Institute of Health); pLAMP1-miRFP703 (Addgene plasmid #79998)^66^ from Dr. Vladislav Verkhusha (Albert Einstein College of Medicine); GFP-C1-PLCdelta-PH (Addgene plasmid #21179)^67^ and Lyn11-targeted FRB (LDR) (Addgene plasmid #20147)^68^ from Dr. Tobias Meyer (Weill Cornell Medical College of Cornell University); mEGFP-C1 (Addgene plasmid #54759) and mTagBFP2-Lifeact-7 (Addgene plasmid #54602)^69^ from Dr. Michael W. Davidson (Florida State University); and CHRM1-Tango (Addgene plasmid #66248)^70^ from Dr. Bryan Roth (The University of North Carolina at Chapel Hill). For the protein expression in *E. coli* cells, pQCSoHis^71^ (provided by Dr. Hiroshi Murakami, Nagoya University) was used as a vector backbone. All expression plasmids were generated using standard cloning procedures and the NEBuilder HiFi DNA assembly system (New England Biolabs). All PCR amplified sequences were verified by DNA sequencing. Complete plasmid sequences are available upon request.

### Protein expression and purification

For *in vitro* experiments, the ORP9-PH domain was expressed as a C-terminal NusA-His6-tag fusion protein (ORP9-PH-NusA-His6) and purified using a method similar to that described previously.^71^ The plasmid encoding the fusion protein (pT5-ORP9-PH-NusA-His6) was transformed into *E. coli.* Strain BL21(DE3) and the transformants were first cultured in 5 mL of LB broth containing 100 μg/mL ampicillin and 5% glucose at 37°C. The culture was transferred to 400 mL of LB broth containing 100 μg/mL ampicillin and 5% glucose, and cells were grown at 37°C until the OD_660_ reached 0.6. Isopropyl-β-D-thiogalactoside was then added to a final concentration of 500 μM to induce protein expression. The cells expressing ORP9-PH-NusA-His6 were further cultured at 18°C for 24 h. The cells were collected by centrifugation, resuspended in buffer A (300 mM AcOK, 1 M KCl, 20 mM HEPES-KOH, 5 mM imidazole, pH 7.8), and disrupted by sonication on ice. The lysate was cleared by centrifugation and purified by a HisTrap FF column (Cytiva) according to the manufacturer’s protocol using a ÄKTA start system (Cytiva). The bound proteins were washed with buffer B (300 mM AcOK, 1 M KCl, 20 mM HEPES-KOH, 10 mM imidazole, 2 mM ATP, 0.2 mM DTT, pH 7.8) and subsequently washed with buffer C (300 mM AcOK, 20 mM HEPES-KOH, 10 mM imidazole, 0.2 mM DTT, pH 7.8) and eluted with buffer D (300 mM AcOK, 500 mM imidazole, 0.2 mM DTT, pH 7.8). The concentration of the protein was determined by UV spectroscopy using the molar extinction coefficient at 280 nm of 55,390 M^-1^ cm^-1^.^72^

### Isothermal titration calorimetry

The binding affinity of ORP9-PH-NusA-His6 to PI4P was determined using a MicroCal iTC200 instrument. All experiments were performed in the same buffer used for dialysis (20 mM HEPES, pH 7.5, 150 mM NaCl). The cell and syringe were filled with 20 µM ORP9-PH-NusA-His6 and 600 µM PI4P (08:0 PI4P, Avanti), respectively. An initial injection of 0.5 µL PI4P was followed by 15 injections of 2.3 µL at time intervals of 2 min at 25°C. The syringe was stirred at 1,000 rpm. The data were analyzed using the ORIGIN software (MicroCal) with a one-site binding model to extract the thermodynamic parameters *K*_a_ (1/*K*_d_), Δ*H*, and the stoichiometry *N*. Δ*G* and Δ*S* were derived from the relations Δ*G* = –RTln*K*_a_ and Δ*G* = Δ*H*–TΔ*S*.

### Cell culture

HeLa, Cos-7, and HEK293 cells were obtained from the Cell Resource Center for Biomedical Research, Institute of Development, Aging and Cancer, Tohoku University. All cells were cultured in Dulbecco’s modified Eagle’s medium (DMEM) supplemented with 10% heat-inactivated fetal bovine serum (FBS), 100 U/mL penicillin, and 100 µg/mL streptomycin at 37°C under a humidified 5% CO_2_ atmosphere.

### Live cell imaging

Confocal fluorescence imaging was performed on an Olympus IX83/FV3000 confocal laser-scanning microscope equipped with a PlanApo N 60×/1.42 NA oil objective (Olympus), a Z drift compensator system (IX3-ZDC2, Olympus), and a stage top incubator (Tokai Hit). Time-lapse live-cell imaging was performed on an IX83 spinning disk confocal microscope (Olympus) equipped with a UPLXAPO60XO/1.42 NA oil objective lens (Olympus), a Z drift compensator system (IX3-ZDC2, Olympus), a CMOS camera (Orca-Flash4.0 V3; Hamamatsu Photonics), a spinning disk confocal unit (CSU-W1; Yokogawa Electric Corp.), and a stage top incubator (Tokai Hit). Fluorescence images were acquired using lasers at 405 nm (for mTagBFP2), 488 nm (for AcGFP1, mEGFP, and AzamiGreen), 561 nm (for dTomato, mScarlet-I, and mCherry), and 640 nm (for iRFP713 and miRFP703) at 37°C under a 5% CO_2_ atmosphere. For all imaging experiments, phenol red- and serum-free DMEM supplemented with 100 U/mL penicillin and 100 µg/mL streptomycin [DMEM(–)] was used. Fluorescence images were analyzed using the Fiji distribution of ImageJ.^73^ For region-of-interest (ROI) analysis, the background fluorescence intensity was subtracted from the total fluorescence intensity.

### Colocalization assays

For colocalization assays, cells were plated at 7 × 10^4^ cells in 35-mm glass-bottomed dishes (IWAKI) and cultured for 24 h at 37°C. The cells were cotransfected with the following plasmids using 293fectin transfection reagent (Thermo Fisher Scientific) according to the manufacturer’s instructions at a 3:7 ratio (0.3 µg and 0.7 µg): (i) pCMV-AcGFP1-ORP9-PH and either of pCMV-GalT-mCherry (Golgi marker), pCMV-LAMP1-mCherry (lysosome marker), piRFP-FRB-Rab7 (Addgene plasmid #51613)^39^ (late endosome marker), or piRFP-FRB-Rab5 (Addgene plasmid #51612)^39^ (early endosome marker); (ii) pCMV-dTomato-ORP9-PH and either of pAcGFP1-Golgi (Golgi marker), pLAMP1-miRFP703 (Addgene plasmid #79998)^66^ (lysosome marker), piRFP-FRB-Rab7 (late endosome marker), or piRFP-FRB-Rab5 (early endosome marker); (iii) pCMV-mTagBFP2-ORP9-PH and either of pCMV-GalT-mCherry (Golgi marker), pCMV-LAMP1-mCherry (lysosome marker), piRFP-FRB-Rab7 (late endosome marker), or piRFP-FRB-Rab5 (early endosome marker); (iv) pCMV-iRFP713-ORP9-PH and either of pCMV-GalT-mCherry (Golgi marker), pCMV-LAMP1-mCherry (lysosome marker), pCMV-mTagBFP2-FRB-Rab7 (late endosome marker), or pCMV-mTagBFP2-FRB-Rab5 (early endosome marker). Twenty-four hours after transfection, the medium was changed to DMEM(–), and the cells were observed by confocal fluorescence imaging.

For investigation of other PI4P reporters, cells were prepared as described above and co-transfected with the following plasmids using 293fectin at a 3:7 ratio (0.3 µg and 0.7 µg): (i) pCMV-AcGFP1-OSBP-PH and pCMV-GalT-mCherry; (ii) pCMV-AcGFP1-P4M and pCMV-GalT-mCherry; (iii) pCMV-mScarlet-I-ORP9-PH and pAcGFP1-Golgi; (iv) pCMV-AzamiGreen-ORP9-PH and pCMV-GalT-mCherry. Twenty-four hours after transfection, the medium was changed to DMEM(–), and the cells were observed by confocal fluorescence imaging.

### Lipid depletion experiments

PM PI4P and PI(4,5)P_2_ depletion experiments were performed as previously described.^44^ For PM PI4P depletion experiments, HeLa cells were plated at 7 × 10^4^ cells in 35-mm glass-bottomed dishes and cultured for 24 h at 37°C. The cells were cotransfected with pCMV-AcGFP1-ORP9-PH and pCMV-mTagBFP2-PLCδ-PH using 293fectin at a 1:1 ratio (each 0.5 µg). Twenty-four hours after transfection, the medium was changed to DMEM(–), and the cells were imaged before and 20 min after treatment with A1 (100 nM) or DMSO (0.01%).

For PM PI(4,5)P_2_ depletion experiments, HeLa cells were prepared as described above and co-transfected with pCMV-AcGFP1-ORP9-PH, pCMV-mTagBFP2-PLCδ-PH, pLyn11-FRB (Addgene plasmid #20147),^68^ and pCMV-mCherry-FKBP-INPP5E^44^ using 293fectin at a 1:1:1:1 ratio (each 0.25 µg). Twenty-four hours after transfection, the medium was changed to DMEM(–), and the cells were imaged before and 1 min after treatment with rapamycin (200 nM) or DMSO (0.1%).

### Visualization of PI4P dynamics upon synthetic ORP5-mediated ER–PM contact manipulation

For this experiment, a HeLa cell line stably expressing AcGFP1-ORP9-PH was established using a piggyBac transposon system.^74^ HeLa cells were cotransfected with pPBbsr-AcGFP1-ORP9-PH and pCMV-mPBase^75^ encoding the piggyBac transposase (provided by Dr. Allan Bradley, Wellcome Trust Sanger Institute) using 293fectin. Cells were selected with 10 μg/mL blasticidin S for at least 10 days. Bulk populations of selected cells were used for the following experiments.

To visualize PI4P dynamics upon synthetic ORP5-mediated ER–PM contact manipulation, HeLa cells stably expressing AcGFP1-ORP9-PH were plated at 7 × 10^4^ cells in 35-mm glass-bottomed dishes and cultured for 24 h at 37°C. The cells were cotransfected with pCMV-mScarlet-I-eDHFR-ORP5(ΔPH) and pCAGGS-HaloTag-KRas4B(CT) using Polyethyleneimine “Max” transfection reagent (PEI-Max) (Polysciences) according to the manufacturer’s instructions at a 3:7 ratio (0.3 µg and 0.7 µg). Twenty-four hours after transfection, the medium was changed to DMEM(–), and the cells were observed by time-lapse imaging (i) before and after the stepwise addition of TMP-HL^56,57^ (0.5 µM) and TMP (100 µM), (ii) before and after the addition of TMP-HL (0.5 µM), or (iii) before and after the addition of DMSO (0.05%).

### Visualization of PI4P dynamics upon M1R stimulation

To visualize PI4P dynamics upon M1R stimulation, HeLa cells were plated at 7 × 10^4^ cells in 35-mm glass-bottomed dishes and cultured for 24 h at 37°C. The cells were cotransfected with pCMV-AcGFP1-ORP9-PH and pCHRM1-Tango (Addgene plasmid #66248)^70^ using PEI-Max at a 1:1 ratio (each 0.5 µg). Twenty-four hours after transfection, the medium was changed to DMEM(–), and the cells were observed by time-lapse imaging (i) before and after the stepwise addition of carbachol (1 mM) (Tokyo Chemical Industry) and atropine (10 µM) (Tokyo Chemical Industry), (ii) before and after the addition of carbachol (1 mM), or (iii) before and after the addition of DMSO (0.1%).

## Acknowledgments

This work was supported by JSPS KAKENHI grant nos. JP20H04706, JP21H02078, and JP21H05226 (S.T.). This work was also supported by JSPS KAKENHI grant nos. JP22H04641 and JP21H02694 (F.N.), the Takeda Science Foundation (F.N.), JSPS KAKENHI grant nos. JP19H05798 and JP22H02625 (K.A.), Joint Research by the National Institutes of Natural Sciences (NINS) (NINS program No. 01112202) (S.T., K.A., F.N.), and Joint Research of the Exploratory Research Center on Life and Living Systems (ExCELLS) (ExCELLS program No. 23EXC601-1) (F.N., K.A., S.T.).

