## Supplementary Information for "ORP9-PH domain-based fluorescent reporters for visualizing phosphatidylinositol 4-phosphate dynamics in living cells"

### Supplementary Figures

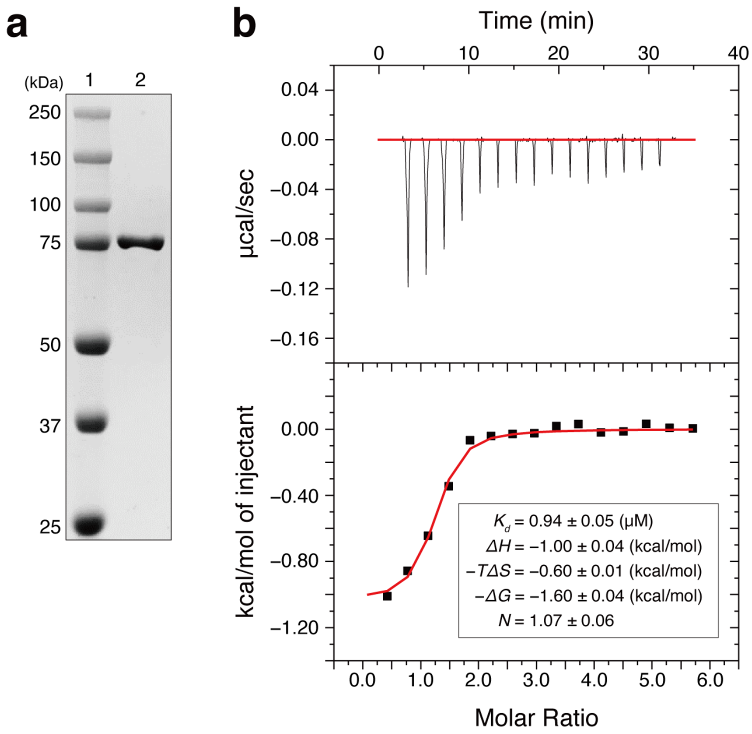


**Figure S1.** Affinity measurement of the ORP9-PH domain to PI4P by ITC. (**a**) SDS-PAGE image of the recombinant ORP9-PH domain used for ITC experiments. Lane 1, molecular weight marker; lane 2, the purified ORP9-PH domain fused to NusA-His6-tag (ORP9-PH-NusA-His6) (calculated M.W.: 72 kDa). (**b**) ITC titration kinetics (top) and integrated binding isotherm (bottom). Measurement conditions: [ORP9-PH-NusA-His6] = 20 μM, [08:0 PI4P] = 600 μM (15 × 2.3 μL injections), 20 mM HEPES buffer, 150 mM NaCl, pH 7.5, 25ºC. Thermodynamic parameters corresponding to the binding of ORP9-PH-NusA-His6 to PI4P are given in the inserted table.

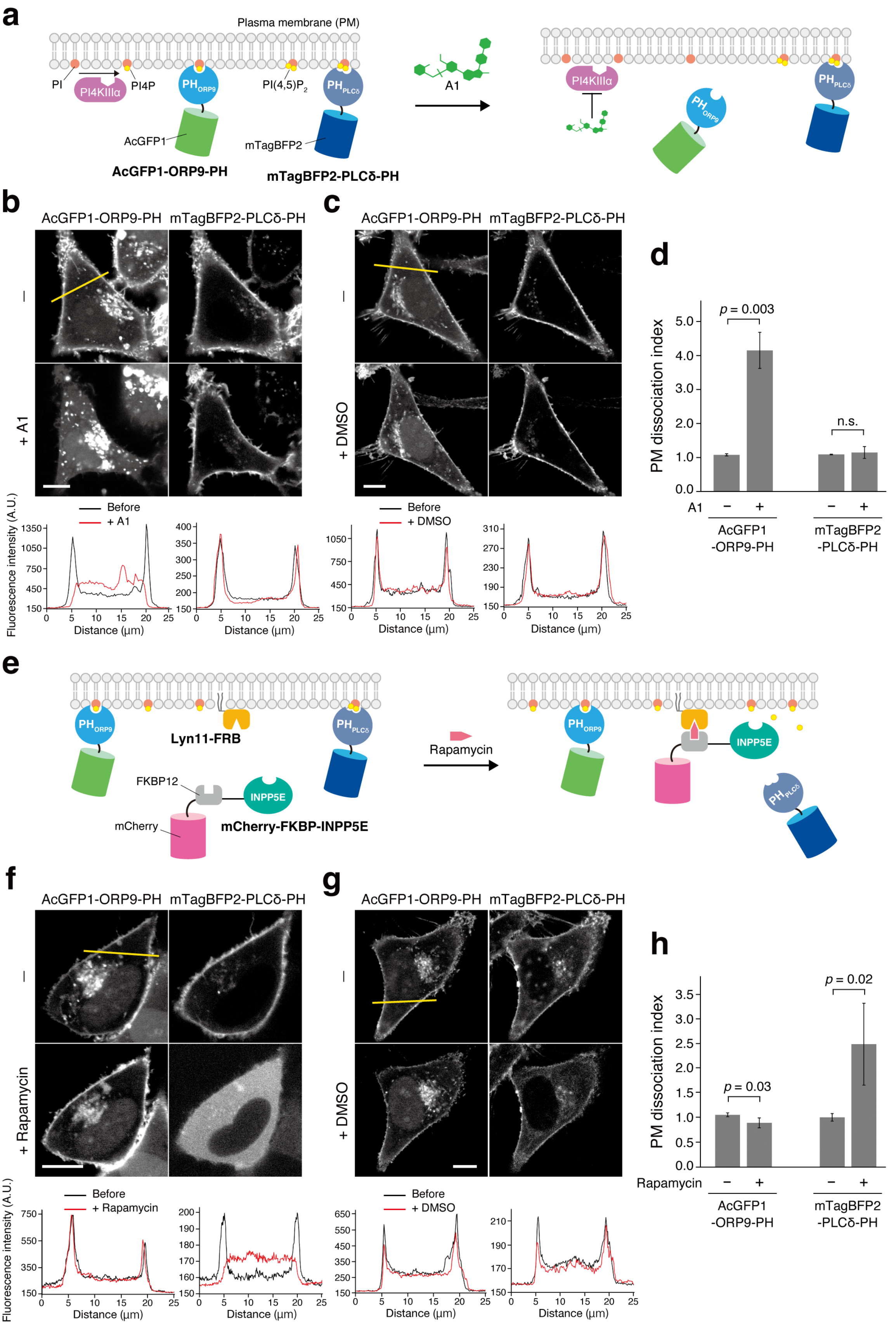


**Figure S2.** Investigation of PI4P specificity of AcGFP1-ORP9-PH. (**a**) Schematic illustration of the PI4P depletion experiment using the PI4KIIIα inhibitor A1. (**b**) Confocal fluorescence images of HeLa cells coexpressing AcGFP1-ORP9-PH (left) and mTagBFP2-PLCδ-PH [PI(4,5)$P_2$ reporter] (right) were taken before (top) and 20 min after the addition of A1 (100 nM) (bottom). Fluorescence intensity profiles of AcGFP1-ORP9-PH and mTagBFP2-PLCδ-PH across the yellow line before (black) and after the A1 addition (red) are shown below the image. Scale bar, 10 μm. (**c**) Control experiment for **b** using DMSO. Scale bar, 10 μm. (**d**) Quantification of PM dissociation of AcGFP1-ORP9-PH and mTagBFP2-PLCδ-PH by PM PI4P depletion. The PM dissociation index is given by $(F_{cyto}/F_{PM})/(F_{cyto}/F_{PM})_0$, where $(F_{cyto}/F_{PM})$ and $(F_{cyto}/F_{PM})_0$ are the ratios of the cytosolic to the PM fluorescence intensity after and before lipid depletion, respectively. Data are presented as the mean ± SD (n = 5 cells). P values indicate the results of Student's *t*-test analysis: n.s., $p > 0.05$. (**e**) Schematic illustration of the PI(4,5)$P_2$ depletion experiment using the rapamycin-induced protein dimerization system. In this experiment, PM PI(4,5)$P_2$ was depleted by recruiting FKBP12-tagged INPP5E (mCherry-FKBP-INPP5E) to the PM-localized Lyn11-FRB by the addition of rapamycin. (**f**) Confocal fluorescence images of HeLa cells coexpressing AcGFP1-ORP9-PH (left) and mTagBFP2-PLCδ-PH [PI(4,5)$P_2$ reporter] (right) were taken before (top) and 1 min after the addition of rapamycin (200 nM) (bottom). Fluorescence intensity profiles of AcGFP1-ORP9-PH and mTagBFP2-PLCδ-PH across the yellow line before (black) and after the rapamycin addition (red) are shown below the image. Scale bar, 10 μm. (**g**) Control experiment for **f** using DMSO. Scale bar, 10 μm. (**h**) Quantification of PM dissociation of AcGFP1-ORP9-PH and mTagBFP2-PLCδ-PH by PM PI(4,5)$P_2$ depletion. The details are as given in panel **d**. Data are presented as the mean ± SD (n = 5 cells).

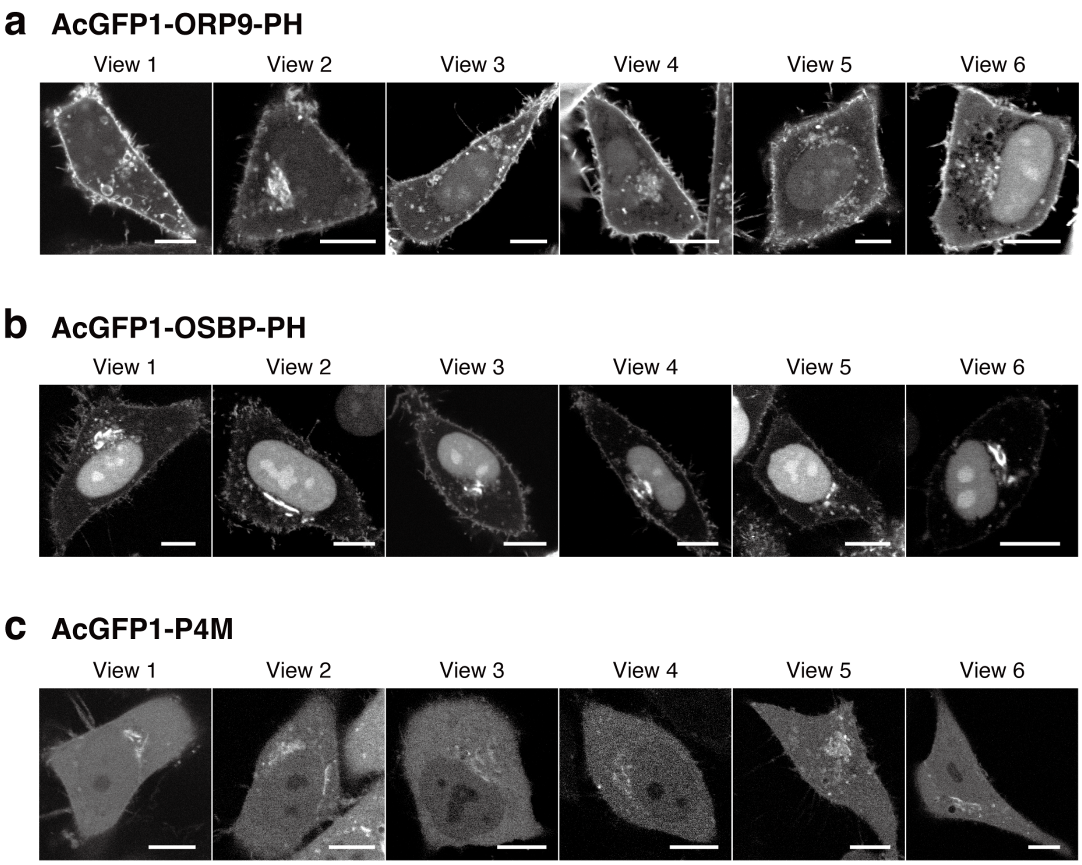


**Figure S3.** Additional confocal fluorescence images of HeLa cells expressing AcGFP1-ORP9-PH (**a**), AcGFP1-OSBP-PH (**b**), and AcGFP1-P4M(**c**), with main images shown in **Fig. 1b**. Scale bars, 10 μm.

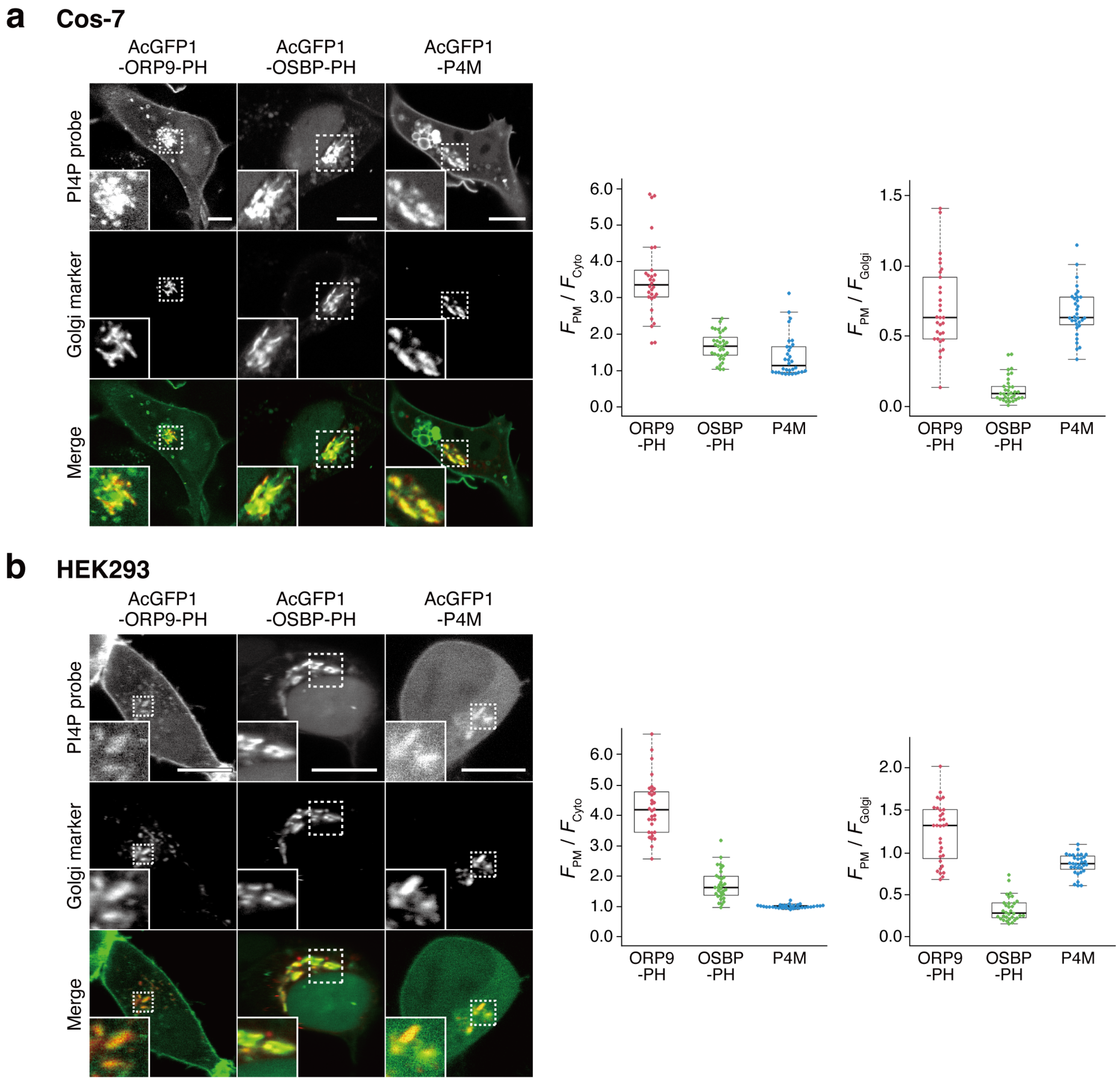


**Figure S4.** Application of fluorescent PI4P reporters (AcGFP1-ORP9-PH, AcGFP1-OSBP-PH, and AcGFP1-P4M) to other cell lines. (**a**) Cos-7 cells. (**b**) HEK293 cells. (Left) Representative confocal fluorescence images of cells coexpressing the indicated PI4P reporter and Golgi marker (GalT-mCherry) are shown. Insets show the regions indicated by white dashed boxes at higher magnification. Scale bars, 10 μm. (Right) Quantification of the PI4P detection sensitivity and Golgi bias. Data analysis was performed as described in **Fig. 1c** ($n \geq 30$ cells).

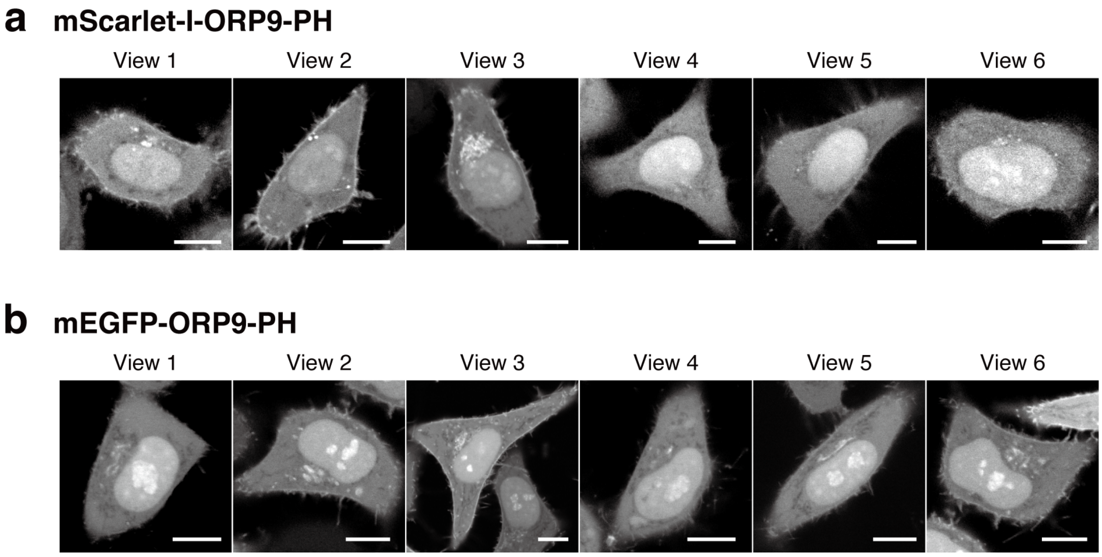


**Figure S5.** Confocal fluorescence images of HeLa cells expressing monomeric fluorescent protein-tagged ORP9-PH domains. (**a**) mScarlet-I-ORP9-PH. (**b**) mEGFP-ORP9-PH. Scale bars, 10 µm.

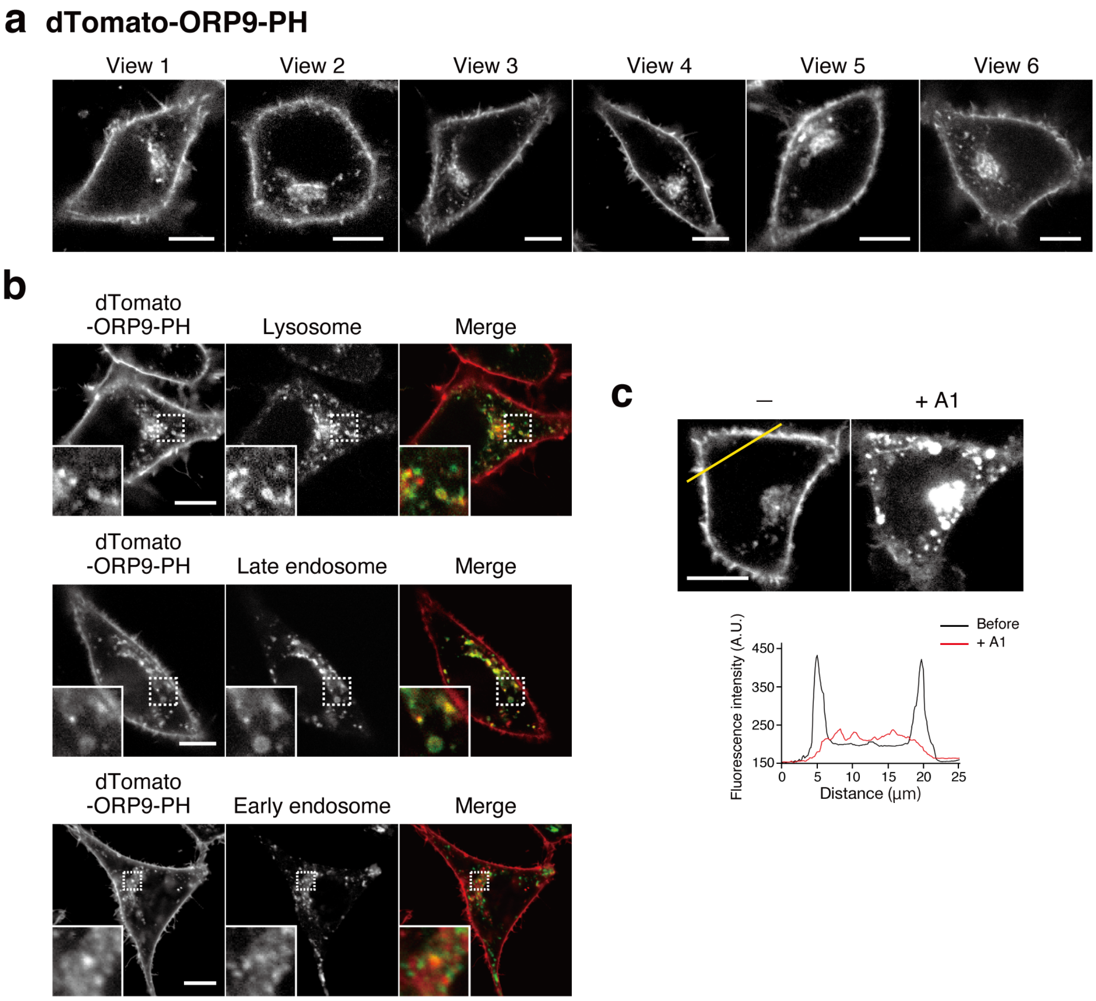


**Figure S6.** PI4P-binding properties of dTomato-ORP9-PH. (**a**) Confocal fluorescence images of HeLa cells expressing dTomato-ORP9-PH. Scale bars, 10 μm. (**b**) Confocal fluorescence images of HeLa cells coexpressing dTomato-ORP9-PH and organelle markers: lysosome, LAMP1-miRFP703; late endosome, iRFP713-Rab7; early endosome: iRFP713-Rab5. Insets show the regions indicated by white dashed boxes at higher magnification. Scale bars, 10 μm. (**c**) PM PI4P depletion experiment. Confocal fluorescence images of HeLa cells expressing dTomato-ORP9-PH were taken before (left) and 20 min after the addition of A1 (100 nM) (right). Fluorescence intensity profiles of dTomato-ORP9-PH across the yellow line before (black) and after the A1 addition (red) are shown below the image. Scale bar, 10 μm.

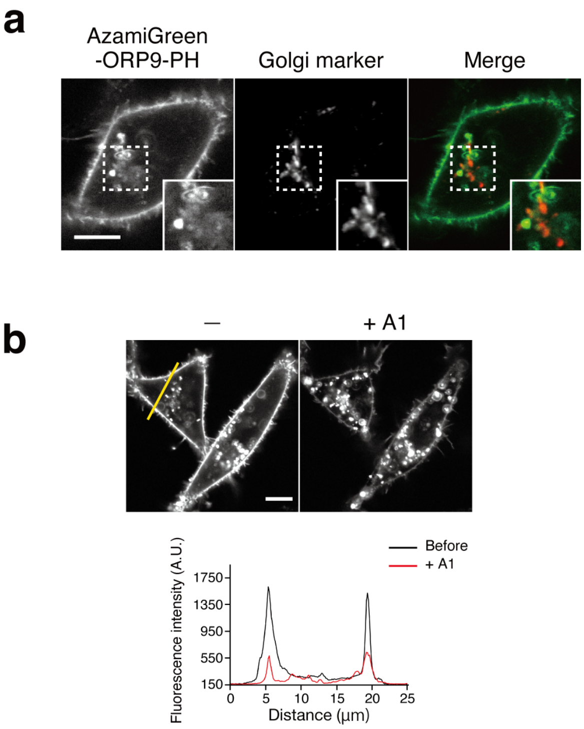


**Figure S7.** Intracellular properties of AzamiGreen-ORP9-PH. (**a**) Confocal fluorescence images of HeLa cells coexpressing AzamiGreen-ORP9-PH and Golgi marker (GalT-mCherry). Insets show the regions indicated by white dashed boxes at higher magnification. Scale bar, 10 μm. (**b**) PM PI4P depletion experiment. Confocal fluorescence images of HeLa cells expressing AzamiGreen-ORP9-PH were taken before (left) and 20 min after the addition of A1 (100 nM) (right). Fluorescence intensity profiles of AzamiGreen-ORP9-PH across the yellow line before (black) and after the A1 addition (red) are shown below the image. Scale bar, 10 μm.

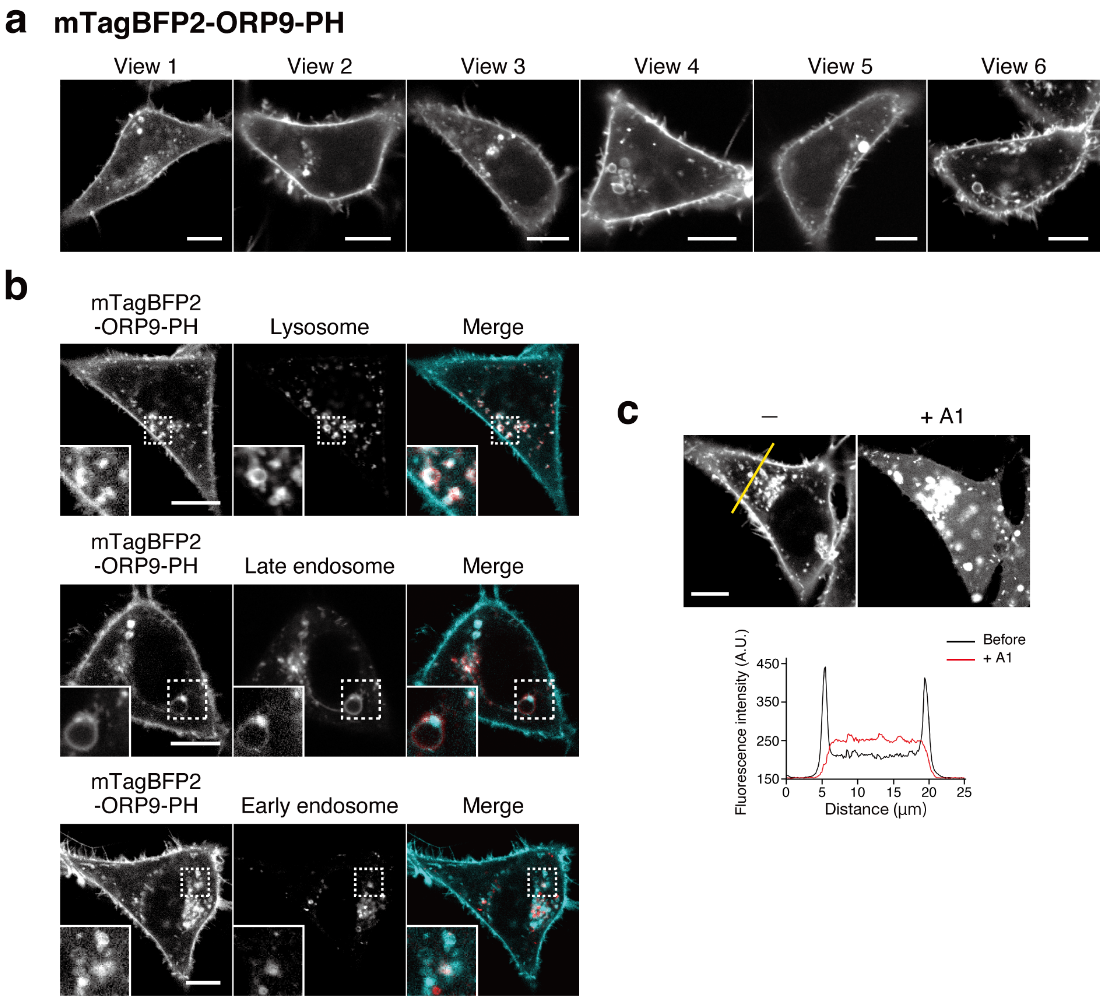


**Figure S8.** PI4P-binding properties of mTagBFP2-ORP9-PH. (**a**) Confocal fluorescence images of HeLa cells expressing mTagBFP2-ORP9-PH. Scale bars, 10 μm. (**b**) Confocal fluorescence images of HeLa cells coexpressing mTagBFP2-ORP9-PH and organelle markers: lysosome, LAMP1-mCherry; late endosome, iRFP713-Rab7; early endosome: iRFP713-Rab5. Insets show the regions indicated by white dashed boxes at higher magnification. Scale bars, 10 μm. (**c**) PM PI4P depletion experiment. Confocal fluorescence images of HeLa cells expressing mTagBFP2-ORP9-PH were taken before (left) and 20 min after the addition of A1 (100 nM) (right). Fluorescence intensity profiles of mTagBFP2-ORP9-PH across the yellow line before (black) and after the A1 addition (red) are shown below the image. Scale bar, 10 μm.

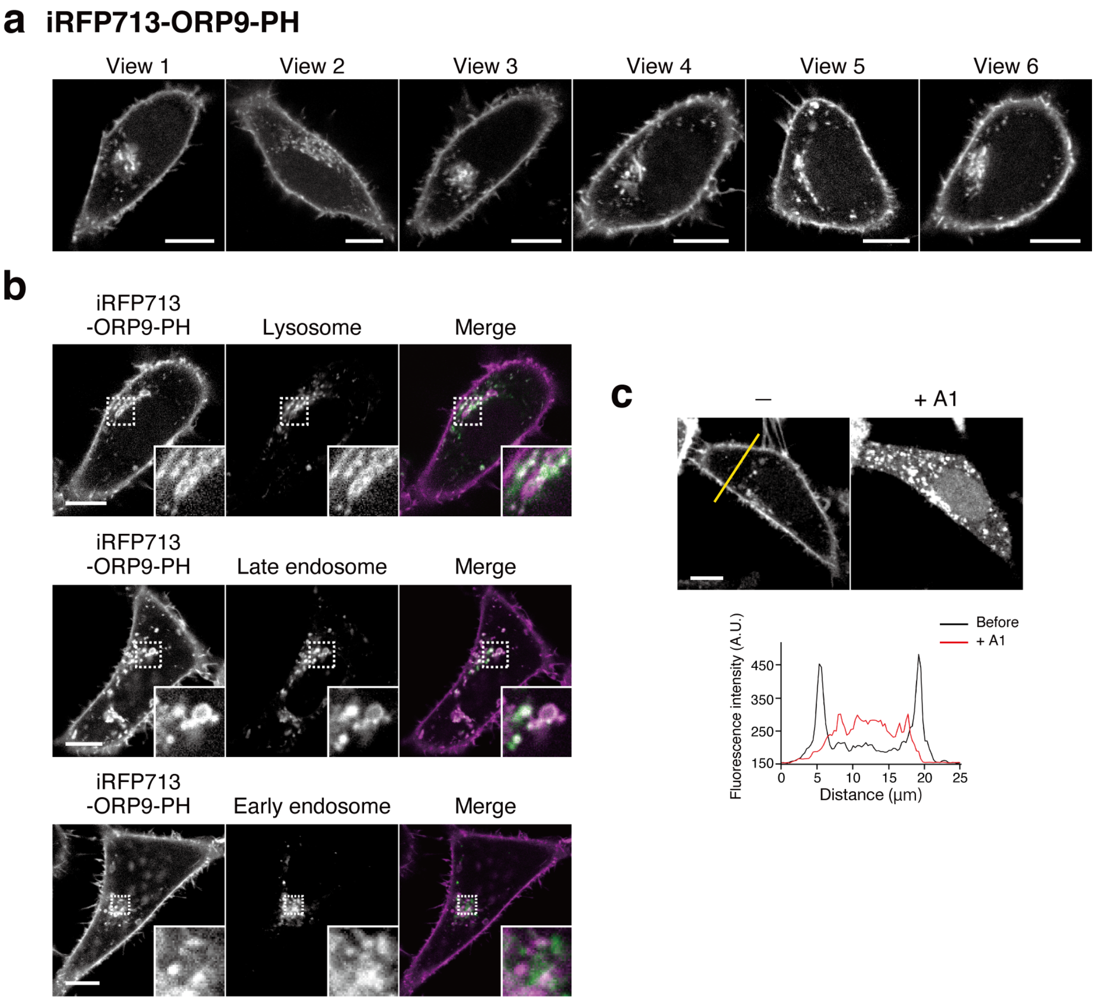


**Figure S9.** PI4P-binding properties of iRFP713-ORP9-PH. (**a**) Confocal fluorescence images of HeLa cells expressing iRFP713-ORP9-PH. Scale bars, 10 µm. (**b**) Confocal fluorescence images of HeLa cells coexpressing iRFP713-ORP9-PH and organelle markers: lysosome, LAMP1-mCherry; late endosome, mTagBFP2-Rab7; early endosome: mTagBFP2-Rab5. Insets show the regions indicated by white dashed boxes at higher magnification. Scale bars, 10 µm. (**c**) PM PI4P depletion experiment. Confocal fluorescence images of HeLa cells expressing iRFP713-ORP9-PH were taken before (left) and 20 min after the addition of A1 (100 nM) (right). Fluorescence intensity profiles of iRFP713-ORP9-PH across the yellow line before (black) and after the A1 addition (red) are shown below the image. Scale bar, 10 µm.

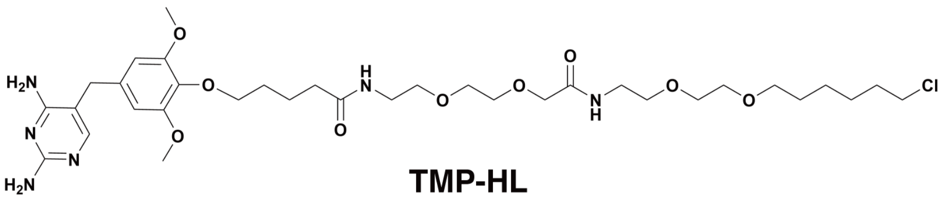

**TMP-HL**

**Figure S10.** Chemical structure of TMP-HL.

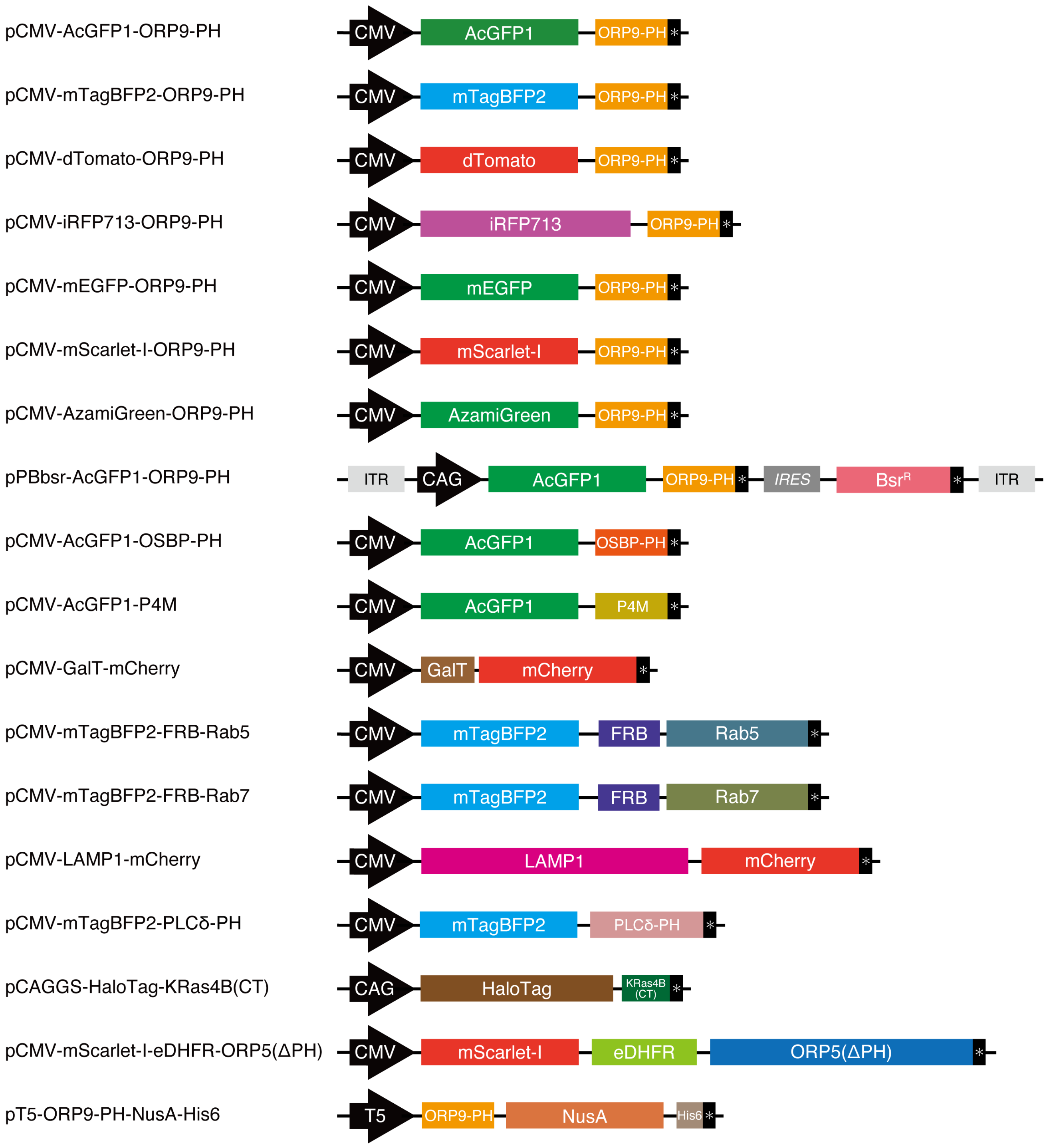


**Figure S11.** Schematic illustration of the domain structures of fusion proteins constructed in this study. DNA and amino acid sequences of the constructs are shown in the "**Supplementary Sequences**" section.

### Supplementary Movies

**Movie S1.** Visualization of PI4P dynamics with AcGFP1-ORP9-PH upon synthetic ORP5-mediated ER–PM contact manipulation (time-lapse movie of **Figure 3b**).

**Movie S2.** Visualization of PM PI4P dynamics with AcGFP1-ORP9-PH upon M1R stimulation (time-lapse movie of **Figure 4a**).

For all movies, scale bars are 10 µm.

### Supplementary Sequences

**pCMV-AcGFP1-ORP9-PH**

>Amino acid sequence
MVSKGAELFTGIVPILIELNGDVNGHKFSVSGEGEGDATYGKLTLKFICTTGKLPVPWPTLVTTLSYGVQCFSRYPDHMKQ
HDFFKSAMPEGYIQERTIFFEDDGNYKSRAEVKFEGDTLVNRIELTGTDFKEDGNILGNKMEYNYNAHNVYIMTDKAKNGI
KVNFKIRHNIEDGSVQLADHYQQNTPIGDGPVLLPDNHYLSTQSALSKDPNEKRDHMIYFGFVTAAAITHGMDELYKSGLR
SGGGGSGGGGSGGGGSRAQASMASIVEGPLSKWTNVMKGWQYRWFVLDYNAGLLSYYTSKDKMMRGSRRGCVRLRGAVIGI
DDEDDSTFTITVDQKTFHFQARDADEREKWIHALEETILRHTLQLQGLDSG*
>DNA sequence
ATGGTGAGCAAGGGCGCCGAGCTGTTCACCGGCATCGTGCCCATCCTGATCGAGCTGAATGGCGATGTGAATGGCCACAAG
TTCAGCGTGAGCGGCGAGGGCGAGGGCGATGCCACCTACGGCAAGCTGACCCTGAAGTTCATCTGCACCACCGGCAAGCTG
CCTGTGCCCTGGCCCACCCTGGTGACCACCCTGAGCTACGGCGTGCAGTGCTTCTCACGCTACCCCGATCACATGAAGCAG
CACGACTTCTTCAAGAGCGCCATGCCTGAGGGCTACATCCAGGAGCGCACCATCTTCTTCGAGGATGACGGCAACTACAAG
TCGCGCGCCGAGGTGAAGTTCGAGGGCGATACCCTGGTGAATCGCATCGAGCTGACCGGCACCGATTTCAAGGAGGATGGC
AACATCCTGGGCAATAAGATGGAGTACAACTACAACGCCCACAATGTGTACATCATGACCGACAAGGCCAAGAATGGCATC
AAGGTGAACTTCAAGATCCGCCACAACATCGAGGATGGCAGCGTGCAGCTGGCCGACCACTACCAGCAGAATACCCCCATC
GGCGATGGCCCTGTGCTGCTGCCCGATAACCACTACCTGTCCACCCAGAGCGCCCTGTCCAAGGACCCCAACGAGAAGCGC
GATCACATGATCTACTTCGGCTTCGTGACCGCCGCCGCCATCACCCACGGCATGGATGAGCTGTACAAGTCCGGACTCAGA
TCTGGTGGCGGAGGCTCGGGCGGAGGTGGGTCGGGTGGCGGCGGATCTCGAGCTCAAGCTTCGATGGCGTCCATCGTGGAA
GGGCCGCTGAGCAAATGGACTAACGTGATGAAGGGATGGCAGTATCGTTGGTTCGTGCTGGACTACAATGCAGGGCTGCTC
TCCTACTACACGTCCAAGGACAAAATGATGAGAGGCTCTCGAAGAGGATGCGTTAGACTCAGAGGAGCTGTGATTGGTATA
GACGACGAGGACGACAGCACCTTCACAATCACTGTCGATCAGAAAACCTTCCACTTCCAGGCTCGAGATGCAGACGAGCGA
GAGAAGTGGATCCATGCCTTAGAAGAAACTATTCTTCGCCATACTCTTCAGCTTCAAGGTTTGGATTCAGGATGA
>Components AcGFP1 ORP9-PH

**pCMV-mTagBFP2-ORP9-PH**

>Amino acid sequence
MVSKGEELIKENMHMKLYMEGTVDNHHFKCTSEGEGKPYEGTQTMRIKVVEGGPLPFAFDILATSFLYGSKTFINHTQGIP
DFFKQSFPEGFTWERVTTYEDGGVLTATQDTSLQDGCLIYNVKIRGVNFTSNGPVMQKKTLGWEAFTETLYPADGGLEGRN
DMALKLVGGSHLIANAKTTYRSKKPAKNLKMPGVYYVDYRLERIKEANNETYVEQHEVAVARYCDLPSKLGHKLNSGLRSG
GGGSGGGGSGGGGSRAQASMASIVEGPLSKWTNVMKGWQYRWFVLDYNAGLLSYYTSKDKMMRGSRRGCVRLRGAVIGIDD
EDDSTFTITVDQKTFHFQARDADEREKWIHALEETILRHTLQLQGLDSG*
>DNA sequence
ATGGTGTCTAAGGGCGAAGAGCTGATTAAGGAGAACATGCACATGAAGCTGTACATGGAGGGCACCGTGGACAACCATCAC
TTCAAGTGCACATCCGAGGGCGAAGGCAAGCCCTACGAGGGCACCCAGACCATGAGAATCAAGGTGGTCGAGGGCGGCCCT
CTCCCCTTCGCCTTCGACATCCTGGCTACTAGCTTCCTCTACGGCAGCAAGACCTTCATCAACCACACCCAGGGCATCCCC
GACTTCTTCAAGCAGTCCTTCCCTGAGGGCTTCACATGGGAGAGAGTCACCACATACGAAGACGGGGGCGTGCTGACCGCT
ACCCAGGACACCAGCCTCCAGGACGGCTGCCTCATCTACAACGTCAAGATCAGAGGGGTGAACTTCACATCCAACGGCCCT
GTGATGCAGAAGAAAACACTCGGCTGGGAGGCCTTCACCGAGACGCTGTACCCCGCTGACGGCGGCCTGGAAGGCAGAAAC
GACATGGCCCTGAAGCTCGTGGGCGGGAGCCATCTGATCGCAAACGCCAAGACCACATATAGATCCAAGAAACCCGCTAAG
AACCTCAAGATGCCTGGCGTCTACTATGTGGACTACAGACTGGAAAGAATCAAGGAGGCCAACAACGAGACCTACGTCGAG
CAGCACGAGGTGGCAGTGGCCAGATACTGCGACCTCCCTAGCAAACTGGGGCACAAGCTTAATTCCGGACTCAGATCTGGT
GGCGGAGGCTCGGGCGGAGGTGGGTCGGGTGGCGGCGGATCTCGAGCTCAAGCTTCGATGGCGTCCATCGTGGAAGGGCCG
CTGAGCAAATGGACTAACGTGATGAAGGGATGGCAGTATCGTTGGTTCGTGCTGGACTACAATGCAGGGCTGCTCTCCTAC
TACACGTCCAAGGACAAAATGATGAGAGGCTCTCGAAGAGGATGCGTTAGACTCAGAGGAGCTGTGATTGGTATAGACGAC
GAGGACGACAGCACCTTCACAATCACTGTCGATCAGAAAACCTTCCACTTCCAGGCTCGAGATGCAGACGAGCGAGAGAAG
TGGATCCATGCCTTAGAAGAAACTATTCTTCGCCATACTCTTCAGCTTCAAGGTTTGGATTCAGGATGA
>Components mTagBFP2 ORP9-PH

**pCMV-dTomato-ORP9-PH**

>Amino acid sequence
MVSKGEEVIKEFMRFKVRMEGSMNGHEFEIEGEGEGRPYEGTQTAKLKVTKGGPLPFAWDILSPQFMYGSKAYVKHPADIP
DYKKLSFPEGFKWERVMNFEDGGLVTVTQDSSLQDGTLIYKVKMRGTNFPPDGPVMQKKTMGWEASTERLYPRDGVLKGEI
HQALKLKDGGHYLVEFKTIYMAKKPVQLPGYYYVDTKLDITSHNEDYTIVEQYERSEGRHHLFLYGMDELYKSGLRSGGGG
SGGGGSGGGGSRAQASMASIVEGPLSKWTNVMKGWQYRWFVLDYNAGLLSYYTSKDKMMRGSRRGCVRLRGAVIGIDDEDD
STFTITVDQKTFHFQARDADEREKWIHALEETILRHTLQLQGLDSG*
>DNA sequence
ATGGTGAGCAAGGGCGAGGAGGTCATCAAAGAGTTCATGCGCTTCAAGGTGCGCATGGAGGGCTCCATGAACGGCCACGAG
TTCGAGATCGAGGGCGAGGGCGAGGGCCGCCCCTACGAGGGCACCCAGACCGCCAAGCTGAAGGTGACCAAGGGCGGCCCC
CTGCCCTTCGCCTGGGACATCCTGTCCCCCCAGTTCATGTACGGCTCCAAGGCGTACGTGAAGCACCCCGCCGACATCCCC
GATTACAAGAAGCTGTCCTTCCCCGAGGGCTTCAAGTGGGAGCGCGTGATGAACTTCGAGGACGGCGGTCTGGTGACCGTG
ACCCAGGACTCCTCCCTGCAGGACGGCACGCTGATCTACAAGGTGAAGATGCGCGGCACCAACTTCCCCCCCGACGGCCCC
GTAATGCAGAAGAAGACCATGGGCTGGGAGGCCTCCACCGAGCGCCTGTACCCCCGCGACGGCGTGCTGAAGGGCGAGATC
CACCAGGCCCTGAAGCTGAAGGACGGCGGCCACTACCTGGTGGAGTTCAAGACCATCTACATGGCCAAGAAGCCCGTGCAA
CTGCCCGGCTACTACTACGTGGACACCAAGCTGGACATCACCTCCCACAACGAGGACTACACCATCGTGGAACAGTACGAG

CGCTCCGAGGGCCGCCACCACCTGTTCCTGTACGGCATGGACGAGCTGTACAAGTCCGGACTCAGATCTGGTGGCGGAGGC
TCGGGCGGAGGTGGGTCGGGTGGCGGCGGATCTCGAGCTCAAGCTTCGATGGCGTCCATCGTGGAAGGGCCGCTGAGCAAA
TGGACTAACGTGATGAAGGGATGGCAGTATCGTTGGTTCGTGCTGGACTACAATGCAGGGCTGCTCTCCTACTACACGTCC
AAGGACAAAATGATGAGAGGCTCTCGAAGAGGATGCGTTAGACTCAGAGGAGCTGTGATTGGTATAGACGACGAGGACGAC
AGCACCTTCACAATCACTGTCGATCAGAAAACCTTCCACTTCCAGGCTCGAGATGCAGACGAGCGAGAGAAGTGGATCCAT
GCCTTAGAAGAAACTATTCTTCGCCATACTCTTCAGCTTCAAGGTTTGGATTCAGGATGA
>Components dTomato ORP9-PH

**pCMV-iRFP713-ORP9-PH**
>Amino acid sequence
MAEGSVARQPDLLTCDDEPIHIPGAIQPHGLLLALAADMTIVAGSDNLPELTGLAIGALIGRSAADVFDSETHNRLTIALA
EPGAAVGAPITVGFTMRKDAGFIGSWHRHDQLIFLELEPPQRDVAEPQAFFRRTNSAIRRLQAAETLESACAAAAQEVRKI
TGFDRVMIYRFASDFSGEVIAEDRCAEVESKLGLHYPASTVPAQARRLYTINPVRIIPDINYRPVPVTPDLNPVTGRPIDL
SFAILRSVSPVHLEFMRNIGMHGTMSISILRGERLWGLIVCHHRTPYYVDLDGRQACELVAQVLAWQIGVMEELYKSGLRS
GGGGSGGGGSGGGGSRAQASMASIVEGPLSKWTNVMKGWQYRWFVLDYNAGLLSYYTSKDKMMRGSRRGCVRLRGAVIGID
DEDDSTFTITVDQKTFHFQARDADEREKWIHALEETILRHTLQLQGLDSG*
>DNA sequence
ATGGCGGAAGGATCTGTCGCCAGGCAGCCTGACCTCTTGACCTGCGACGATGAGCCGATCCATATCCCCGGTGCCATCCAA
CCGCATGGACTGCTGCTCGCCCTCGCCGCCGACATGACGATCGTTGCCGGCAGCGACAACCTTCCCGAACTCACCGGACTG
GCGATCGGCGCCCTGATCGGCCGCTCTGCGGCCGATGTCTTCGACTCGGAGACGCACAACCGTCTGACGATCGCCTTGGCC
GAGCCCGGGGCGGCCGTCGGAGCACCGATCACTGTCGGCTTCACGATGCGAAAGGACGCAGGCTTCATCGGCTCCTGGCAT
CGCCATGATCAGCTCATCTTCCTCGAGCTCGAGCCTCCCCAGCGGGACGTCGCCGAGCCGCAGGCGTTCTTCCGCCGCACC
AACAGCGCCATCCGCCGCCTGCAGGCCGCCGAAACCTTGGAAAGCGCCTGCGCCGCCGCGGCGCAAGAGGTGCGGAAGATT
ACCGGCTTCGATCGGGTGATGATCTATCGCTTCGCCTCCGACTTCAGCGGCGAAGTGATCGCAGAGGATCGGTGCGCCGAG
GTCGAGTCAAAACTAGGCCTGCACTATCCTGCCTCAACCGTGCCGGCGCAGGCCCGTCGGCTCTATACCATCAACCCGGTA
CGGATCATTCCCGATATCAATTATCGGCCGGTGCCGGTCACCCCAGACCTCAATCCGGTCACCGGGCGGCCGATTGATCTT
AGCTTCGCCATCCTGCGCAGCGTCTCGCCCGTCCATCTGGAGTTCATGCGCAACATAGGCATGCACGGCACGATGTCGATC
TCGATTTTGCGCGGCGAGCGACTGTGGGGATTGATCGTTTGCCATCACCGAACGCCGTACTACGTCGATCTCGATGGCCGC
CAAGCCTGCGAGCTAGTCGCCCAGGTTCTGGCCTGGCAGATCGGCGTGATGGAAGAGCTGTACAAGTCCGGACTCAGATCT
GGTGGCGGAGGCTCGGGCGGAGGTGGGTCGGGTGGCGGCGGATCTCGAGCTCAAGCTTCGATGGCGTCCATCGTGGAAGGG
CCGCTGAGCAAATGGACTAACGTGATGAAGGGATGGCAGTATCGTTGGTTCGTGCTGGACTACAATGCAGGGCTGCTCTCC
TACTACACGTCCAAGGACAAAATGATGAGAGGCTCTCGAAGAGGATGCGTTAGACTCAGAGGAGCTGTGATTGGTATAGAC
GACGAGGACGACAGCACCTTCACAATCACTGTCGATCAGAAAACCTTCCACTTCCAGGCTCGAGATGCAGACGAGCGAGAG
AAGTGGATCCATGCCTTAGAAGAAACTATTCTTCGCCATACTCTTCAGCTTCAAGGTTTGGATTCAGGATGA
>Components iRFP713 ORP9-PH

**pCMV-mEGFP-ORP9-PH**
>Amino acid sequence
MVSKGEELFTGVVPILVELDGDVNGHKFSVSGEGEGDATYGKLTLKFICTTGKLPVPWPTLVTTLTYGVQCFSRYPDHMKQ
HDFFKSAMPEGYVQERTIFFKDDGNYKTRAEVKFEGDTLVNRIELKGIDFKEDGNILGHKLEYNYNSHNVYIMADKQKNGI
KVNFKIRHNIEDGSVQLADHYQQNTPIGDGPVLLPDNHYLSTQSKLSKDPNEKRDHMVLLEFVTAAGITLGMDELYKSGLR
SGGGGSGGGGSGGGGSRAQASMASIVEGPLSKWTNVMKGWQYRWFVLDYNAGLLSYYTSKDKMMRGSRRGCVRLRGAVIGI
DDEDDSTFTITVDQKTFHFQARDADEREKWIHALEETILRHTLQLQGLDSG*
>DNA sequence
ATGGTGAGCAAGGGCGAGGAGCTGTTCACCGGGGTGGTGCCCATCCTGGTCGAGCTGGACGGCGACGTAAACGGCCACAAG
TTCAGCGTGTCCGGCGAGGGCGAGGGCGATGCCACCTACGGCAAGCTGACCCTGAAGTTCATCTGCACCACCGGCAAGCTG
CCCGTGCCCTGGCCCACCCTCGTGACCACCCTGACCTACGGCGTGCAGTGCTTCAGCCGCTACCCCGACCACATGAAGCAG
CACGACTTCTTCAAGTCCGCCATGCCCGAAGGCTACGTCCAGGAGCGCACCATCTTCTTCAAGGACGACGGCAACTACAAG
ACCCGCGCCGAGGTGAAGTTCGAGGGCGACACCCTGGTGAACCGCATCGAGCTGAAGGGCATCGACTTCAAGGAGGACGGC
AACATCCTGGGGCACAAGCTGGAGTACAACTACAACAGCCACAACGTCTATATCATGGCCGACAAGCAGAAGAACGGCATC
AAGGTGAACTTCAAGATCCGCCACAACATCGAGGACGGCAGCGTGCAGCTCGCCGACCACTACCAGCAGAACACCCCCATC
GGCGACGGCCCCGTGCTGCTGCCCGACAACCACTACCTGAGCACCCAGTCCAAGCTGAGCAAAGACCCCAACGAGAAGCGC
GATCACATGGTCCTGCTGGAGTTCGTGACCGCCGCCGGGATCACTCTCGGCATGGACGAGCTGTACAAGTCCGGACTCAGA
TCTGGTGGCGGAGGCTCGGGCGGAGGTGGGTCGGGTGGCGGCGGATCTCGAGCTCAAGCTTCGATGGCGTCCATCGTGGAA
GGGCCGCTGAGCAAATGGACTAACGTGATGAAGGGATGGCAGTATCGTTGGTTCGTGCTGGACTACAATGCAGGGCTGCTC
TCCTACTACACGTCCAAGGACAAAATGATGAGAGGCTCTCGAAGAGGATGCGTTAGACTCAGAGGAGCTGTGATTGGTATA
GACGACGAGGACGACAGCACCTTCACAATCACTGTCGATCAGAAAACCTTCCACTTCCAGGCTCGAGATGCAGACGAGCGA
GAGAAGTGGATCCATGCCTTAGAAGAAACTATTCTTCGCCATACTCTTCAGCTTCAAGGTTTGGATTCAGGATGA
>Components mEGFP ORP9-PH

**pCMV-mScarlet-I-ORP9-PH**
>Amino acid sequence
MVSKGEAVIKEFMRFKVHMEGSMNGHEFEIEGEGEGRPYEGTQTAKLKVTKGGPLPFSWDILSPQFMYGSRAFIKHPADIP
DYYKQSFPEGFKWERVMNFEDGGAVTVTQDTSLEDGTLIYKVKLRGTNFPPDGPVMQKKTMGWEASTERLYPEDGVLKGDI
KMALRLKDGGRYLADFKTTYKAKKPVQMPGAYNVDRKLDITSHNEDYTVVEQYERSEGRHSTGGMDELYKSGLRSGGGGSG
GGGSGGGGSRAQASMASIVEGPLSKWTNVMKGWQYRWFVLDYNAGLLSYYTSKDKMMRGSRRGCVRLRGAVIGIDDEDDST

FTITVDQKTFHFQARDADEREKWIHALEETILRHTLQLQGLDSG*
>DNA sequence
ATGGTGAGCAAGGGCGAGGCAGTGATCAAGGAGTTCATGCGGTTCAAGGTGCACATGGAGGGCTCCATGAACGGCCACGAG
TTCGAGATCGAGGGCGAGGGCGAGGGCCGCCCCTACGAGGGCACCCAGACCGCCAAGCTGAAGGTGACCAAGGGTGGCCCC
CTGCCCTTCTCCTGGGACATCCTGTCCCCTCAGTTCATGTACGGCTCCAGGGCCTTCATCAAGCACCCCGCCGACATCCCC
GACTACTATAAGCAGTCCTTCCCCGAGGGCTTCAAGTGGGAGCGCGTGATGAACTTCGAGGACGGCGGCGCCGTGACCGTG
ACCCAGGACACCTCCCTGGAGGACGGCACCCTGATCTACAAGGTGAAGCTCCGCGGCACCAACTTCCCTCCTGACGGCCCC
GTAATGCAGAAGAAGACAATGGGCTGGGAAGCGTCCACCGAGCGGTTGTACCCCGAGGACGGCGTGCTGAAGGGCGACATT
AAGATGGCCCTGCGCCTGAAGGACGGCGGCCGCTACCTGGCGGACTTCAAGACCACCTACAAGGCCAAGAAGCCCGTGCAG
ATGCCCGGCGCCTACAACGTCGACCGCAAGTTGGACATCACCTCCCACAACGAGGACTACACCGTGGTGGAACAGTACGAA
CGCTCCGAGGGCCGCCACTCCACCGGCGGCATGGACGAGCTGTACAAGTCCGGACTCAGATCTGGTGGCGGAGGCTCGGGC
GGAGGTGGGTCGGGTGGCGGCGGATCTCGAGCTCAAGCTTCGATGGCGTCCATCGTGGAAGGGCCGCTGAGCAAATGGACT
AACGTGATGAAGGGATGGCAGTATCGTTGGTTCGTGCTGGACTACAATGCAGGGCTGCTCTCCTACTACACGTCCAAGGAC
AAAATGATGAGAGGCTCTCGAAGAGGATGCGTTAGACTCAGAGGAGCTGTGATTGGTATAGACGACGAGGACGACAGCACC
TTCACAATCACTGTCGATCAGAAAACCTTCCACTTCCAGGCTCGAGATGCAGACGAGCGAGAGAAGTGGATCCATGCCTTA
GAAGAAACTATTCTTCGCCATACTCTTCAGCTTCAAGGTTTGGATTCAGGATGA
>Components mScarlet-I ORP9-PH

**pCMV-AzamiGreen-ORP9-PH**
>Amino acid sequence
MVSVIKPEMKIKLCMRGTVNGHNFVIEGEGKGNPYEGTQILDLNVTEGAPLPFAYDILTTVFQYGNRAFTKYPADIQDYFK
QTFPEGYHWERSMTYEDQGICTATSNISMRGDCFFYDIRFDGVNFPPNGPVMQKKTLKWEPSTEKMYVRDGVLKGDVNMAL
LLEGGGHYRCDFKTTYKAKKDVRLPDYHFVDHRIEILKHDKDYNKVKLYENAVARYSMLPSQAKSGLRSGGGGSGGGGSGG
GGSRAQASMASIVEGPLSKWTNVMKGWQYRWFVLDYNAGLLSYYTSKDKMMRGSRRGCVRLRGAVIGIDDEDDSTFTITVD
QKTFHFQARDADEREKWIHALEETILRHTLQLQGLDSG*
>DNA sequence
ATGGTGAGCGTGATCAAGCCCGAGATGAAGATCAAGCTGTGCATGAGGGGCACCGTGAACGGCCACAACTTCGTGATCGAG
GGCGAGGGCAAGGGCAACCCCTACGAGGGCACCCAGATCCTGGACCTGAACGTGACCGAGGGCGCCCCCCTGCCCTTCGCC
TACGACATCCTGACCACCGTGTTCCAGTACGGCAACAGGGCCTTCACCAAGTACCCCGCCGACATCCAGGACTACTTCAAG
CAGACCTTCCCCGAGGGCTACCACTGGGAGAGGAGCATGACCTACGAGGACCAGGGCATCTGCACCGCCACCAGCAACATC
AGCATGAGGGGCGACTGCTTCTTCTACGACATCAGGTTCGACGGCGTGAACTTCCCCCCCAACGGCCCCGTGATGCAGAAG
AAGACCCTGAAGTGGGAGCCCAGCACCGAGAAGATGTACGTGAGGGACGGCGTGCTGAAGGGCGACGTGAACATGGCCCTG
CTGCTGGAGGGCGGCGGCCACTACAGGTGCGACTTCAAGACCACCTACAAGGCCAAGAAGGACGTGAGGCTGCCCGACTAC
CACTTCGTGGACCACAGGATCGAGATCCTGAAGCACGACAAGGACTACAACAAGGTGAAGCTGTACGAGAACGCCGTGGCC
AGGTACAGCATGCTGCCCAGCCAGGCCAAGTCCGGACTCAGATCTGGTGGCGGAGGCTCGGGCGGAGGTGGGTCGGGTGGC
GGCGGATCTCGAGCTCAAGCTTCGATGGCGTCCATCGTGGAAGGGCCGCTGAGCAAATGGACTAACGTGATGAAGGGATGG
CAGTATCGTTGGTTCGTGCTGGACTACAATGCAGGGCTGCTCTCCTACTACACGTCCAAGGACAAAATGATGAGAGGCTCT
CGAAGAGGATGCGTTAGACTCAGAGGAGCTGTGATTGGTATAGACGACGAGGACGACAGCACCTTCACAATCACTGTCGAT
CAGAAAACCTTCCACTTCCAGGCTCGAGATGCAGACGAGCGAGAGAAGTGGATCCATGCCTTAGAAGAAACTATTCTTCGC
CATACTCTTCAGCTTCAAGGTTTGGATTCAGGATGA
>Components AzamiGreen ORP9-PH

**pPBbsr-AcGFP1-ORP9-PH**
>Amino acid sequence
MVSKGAELFTGIVPILIELNGDVNGHKFSVSGEGEGDATYGKLTLKFICTTGKLPVPWPTLVTTLSYGVQCFSRYPDHMKQ
HDFFKSAMPEGYIQERTIFFEDDGNYKSRAEVKFEGDTLVNRIELTGTDFKEDGNILGNKMEYNYNAHNVYIMTDKAKNGI
KVNFKIRHNIEDGSVQLADHYQQNTPIGDGPVLLPDNHYLSTQSALSKDPNEKRDHMIYFGFVTAAAITHGMDELYKSGLR
SGGGGSGGGGSGGGGSRAQASASIVEGPLSKWTNVMKGWQYRWFVLDYNAGLLSYYTSKDKMMRGSRRGCVRLRGAVIGID
DEDDSTFTITVDQKTFHFQARDADEREKWIHALEETILRHTLQLQGLDSGMASIVEGPLSKWTNVMKGWQYRWFVLDYNAG
LLSYYTSKDKMMRGSRRGCVRLRGAVIGIDDEDDSTFTITVDQKTFHFQARDADEREKWIHALEETILRHTLQLQGLDSG*
-[*IRES*]-MLYEDNKHHVGAAIRTKTGEIISAVHIEAYIGRVTVCAEAIAIGSAVSNGQKDFDTIVAVRHPYSDEVDRSIR
VVSPCGMCRELISDYAPDCFVLIEMNGKLVKTTIEELIPLKYTRN*
>DNA sequence
ATGGTGAGCAAGGGCGCCGAGCTGTTCACCGGCATCGTGCCCATCCTGATCGAGCTGAATGGCGATGTGAATGGCCACAAG
TTCAGCGTGAGCGGCGAGGGCGAGGGCGATGCCACCTACGGCAAGCTGACCCTGAAGTTCATCTGCACCACCGGCAAGCTG
CCTGTGCCCTGGCCCACCCTGGTGACCACCCTGAGCTACGGCGTGCAGTGCTTCTCACGCTACCCCGATCACATGAAGCAG
CACGACTTCTTCAAGAGCGCCATGCCTGAGGGCTACATCCAGGAGCGCACCATCTTCTTCGAGGATGACGGCAACTACAAG
TCGCGCGCCGAGGTGAAGTTCGAGGGCGATACCCTGGTGAATCGCATCGAGCTGACCGGCACCGATTTCAAGGAGGATGGC
AACATCCTGGGCAATAAGATGGAGTACAACTACAACGCCCACAATGTGTACATCATGACCGACAAGGCCAAGAATGGCATC
AAGGTGAACTTCAAGATCCGCCACAACATCGAGGATGGCAGCGTGCAGCTGGCCGACCACTACCAGCAGAATACCCCCATC
GGCGATGGCCCTGTGCTGCTGCCCGATAACCACTACCTGTCCACCCAGAGCGCCCTGTCCAAGGACCCCAACGAGAAGCGC
GATCACATGATCTACTTCGGCTTCGTGACCGCCGCCGCCATCACCCACGGCATGGATGAGCTGTACAAGTCCGGACTCAGA
TCTGGTGGCGGAGGCTCGGGCGGAGGTGGGTCGGGTGGCGGCGGATCTCGAGCTCAAGCTTCGGGCTCGGCTCGAGAGGGC
TGGCTCTTCAAATGGACCAATTATATCAAAGGCTACCAGCGGCGATGGTTCGTGCTGAGCAACGGGCTCCTGAGCTACTAC
AGATCAAAGGCAGAGATGAGACATACCTGCCGTGGTACCATCAACCTCGCCACAGCCAACATCACCGTGGAGGACTCCTGC
AACTTCATCATTTCCAATGGGGGTGCTCAGACCTACCATCTGAAAGCTAGTTCAGAAGTTGAGCGGCAGCGCTGGGTGACG
GCCCTGGAACTGGCCAAGGCCAAAGCTGTGAAGTGACCCGGGATCCTGCAGTCGACGGGCCGCGGTAACAATTGTTAACTA

ACTTAAGCTAGCAACGGTTTCCCTCTAGCGGGATCAATTCCGCCCCCCCCCCCTAACGTTACTGGCCGAAGCCGCTTGGAA
TAAGGCCGGTGTGCGTTTGTCTATATGTTATTTTCCACCATATTGCCGTCTTTTGGCAATGTGAGGGCCCGGAAACCTGGC
CCTGTCTTCTTGACGAGCATTCCTAGGGGTCTTTCCCCTCTCGCCAAAGGAATGCAAGGTCTGTTGAATGTCGTGAAGGAA
GCAGTTCCTCTGGAAGCTTCTTGAAGACAAACAACGTCTGTAGCGACCCTTTGCAGGCAGCGGAACCCCCCACCTGGCGAC
AGGTGCCTCTGCGGCCAAAAGCCACGTGTATAAGATACACCTGCAAAGGCGGCACAACCCCAGTGCCACGTTGTGAGTTGG
ATAGTTGTGGAAAGAGTCAAATGGCTCTCCTCAAGCGTATTCAACAAGGGGCTGAAGGATGCCCAGAAGGTACCCCATTGT
ATGGGATCTGATCTGGGGCCTCGGTGCACATGCTTTACATGTGTTTAGTCGAGGTTAAAAAACGTCTAGGCCCCCCGAACC
ACGGGGACGTGGTTTTCCTTTGAAAAACACGATAATACCATGGTCATGAAAACATTTAACATTTCTCAACAAGATCTAGAA
TTAGTAGAAGTAGCGACAGAGAAGATTACAATGCTTTATGAGGATAATAAACATCATGTGGGAGCGGCAATTCGTACGAAA
ACAGGAGAAATCATTTCGGCAGTACATATTGAAGCGTATATAGGACGAGTAACTGTTTGTGCAGAAGCCATTGCGATTGGT
AGTGCAGTTTCGAATGGACAAAAGGATTTTGACACGATTGTAGCTGTTAGACACCCTTATTCTGACGAAGTAGATAGAAGT
ATTCGAGTGGTAAGTCCTTGTGGTATGTGTAGGGAGTTGATTTCAGACTATGCACCAGATTGTTTTGTGTTAATAGAAATG
AATGGCAAGTTAGTCAAAACTACGATTGAAGAACTCATTCCACTCAAATATACCCGAAATTAA
>Components AcGFP1 ORP9-PH *IRES* Bsr$^{R}$(blasticidin S-deaminase)

**pCMV-AcGFP1-OSBP-PH**
>Amino acid sequence
MVSKGAELFTGIVPILIELNGDVNGHKFSVSGEGEGDATYGKLTLKFICTTGKLPVPWPTLVTTLSYGVQCFSRYPDHMKQ
HDFFKSAMPEGYIQERTIFFEDDGNYKSRAEVKFEGDTLVNRIELTGTDFKEDGNILGNKMEYNYNAHNVYIMTDKAKNGI
KVNFKIRHNIEDGSVQLADHYQQNTPIGDGPVLLPDNHYLSTQSALSKDPNEKRDHMIYFGFVTAAAITHGMDELYKSGLR
SGGGGSGGGGSGGGGSRAQASGSAREGWLFKWTNYIKGYQRRWFVLSNGLLSYYRSKAEMRHTCRGTINLATANITVEDSC
NFIISNGGAQTYHLKASSEVERQRWVTALELAKAKAVK*
>DNA sequence
ATGGTGAGCAAGGGCGCCGAGCTGTTCACCGGCATCGTGCCCATCCTGATCGAGCTGAATGGCGATGTGAATGGCCACAAG
TTCAGCGTGAGCGGCGAGGGCGAGGGCGATGCCACCTACGGCAAGCTGACCCTGAAGTTCATCTGCACCACCGGCAAGCTG
CCTGTGCCCTGGCCCACCCTGGTGACCACCCTGAGCTACGGCGTGCAGTGCTTCTCACGCTACCCCGATCACATGAAGCAG
CACGACTTCTTCAAGAGCGCCATGCCTGAGGGCTACATCCAGGAGCGCACCATCTTCTTCGAGGATGACGGCAACTACAAG
TCGCGCGCCGAGGTGAAGTTCGAGGGCGATACCCTGGTGAATCGCATCGAGCTGACCGGCACCGATTTCAAGGAGGATGGC
AACATCCTGGGCAATAAGATGGAGTACAACTACAACGCCCACAATGTGTACATCATGACCGACAAGGCCAAGAATGGCATC
AAGGTGAACTTCAAGATCCGCCACAACATCGAGGATGGCAGCGTGCAGCTGGCCGACCACTACCAGCAGAATACCCCCATC
GGCGATGGCCCTGTGCTGCTGCCCGATAACCACTACCTGTCCACCCAGAGCGCCCTGTCCAAGGACCCCAACGAGAAGCGC
GATCACATGATCTACTTCGGCTTCGTGACCGCCGCCGCCATCACCCACGGCATGGATGAGCTGTACAAGTCCGGACTCAGA
TCTGGTGGCGGAGGCTCGGGCGGAGGTGGGTCGGGTGGCGGCGGATCTCGAGCTCAAGCTTCGGGCTCGGCTCGAGAGGGC
TGGCTCTTCAAATGGACCAATTATATCAAAGGCTACCAGCGGCGATGGTTCGTGCTGAGCAACGGGCTCCTGAGCTACTAC
AGATCAAAGGCAGAGATGAGACATACCTGCCGTGGTACCATCAACCTCGCCACAGCCAACATCACCGTGGAGGACTCCTGC
AACTTCATCATTTCCAATGGGGGTGCTCAGACCTACCATCTGAAAGCTAGTTCAGAAGTTGAGCGGCAGCGCTGGGTGACG
GCCCTGGAACTGGCCAAGGCCAAAGCTGTGAAGTGA
>Components AcGFP1 OSBP-PH

**pCMV-AcGFP1-P4M**
>Amino acid sequence
MVSKGAELFTGIVPILIELNGDVNGHKFSVSGEGEGDATYGKLTLKFICTTGKLPVPWPTLVTTLSYGVQCFSRYPDHMKQ
HDFFKSAMPEGYIQERTIFFEDDGNYKSRAEVKFEGDTLVNRIELTGTDFKEDGNILGNKMEYNYNAHNVYIMTDKAKNGI
KVNFKIRHNIEDGSVQLADHYQQNTPIGDGPVLLPDNHYLSTQSALSKDPNEKRDHMIYFGFVTAAAITHGMDELYKSGLR
STASTENFKNVKEKYQQMRGDALKTEILADFKDKLAEATDEQSLKQIVAELKSKDEYRILAKGQGLTTQLLGLKTSSVSSF
EKMVEETRESIKSQERQTIKIK*
>DNA sequence
ATGGTGAGCAAGGGCGCCGAGCTGTTCACCGGCATCGTGCCCATCCTGATCGAGCTGAATGGCGATGTGAATGGCCACAAG
TTCAGCGTGAGCGGCGAGGGCGAGGGCGATGCCACCTACGGCAAGCTGACCCTGAAGTTCATCTGCACCACCGGCAAGCTG
CCTGTGCCCTGGCCCACCCTGGTGACCACCCTGAGCTACGGCGTGCAGTGCTTCTCACGCTACCCCGATCACATGAAGCAG
CACGACTTCTTCAAGAGCGCCATGCCTGAGGGCTACATCCAGGAGCGCACCATCTTCTTCGAGGATGACGGCAACTACAAG
TCGCGCGCCGAGGTGAAGTTCGAGGGCGATACCCTGGTGAATCGCATCGAGCTGACCGGCACCGATTTCAAGGAGGATGGC
AACATCCTGGGCAATAAGATGGAGTACAACTACAACGCCCACAATGTGTACATCATGACCGACAAGGCCAAGAATGGCATC
AAGGTGAACTTCAAGATCCGCCACAACATCGAGGATGGCAGCGTGCAGCTGGCCGACCACTACCAGCAGAATACCCCCATC
GGCGATGGCCCTGTGCTGCTGCCCGATAACCACTACCTGTCCACCCAGAGCGCCCTGTCCAAGGACCCCAACGAGAAGCGC
GATCACATGATCTACTTCGGCTTCGTGACCGCCGCCGCCATCACCCACGGCATGGATGAGCTGTACAAGTCCGGACTCAGA
TCTACGGCAAGCACGGAAAACTTTAAAAATGTTAAAGAAAAATATCAGCAAATGCGAGGTGATGCTTTAAAAACAGAAATC
CTGGCTGATTTCAAGGATAAACTGGCTGAAGCTACCGATGAACAGAGCCTTAAGCAAATTGTTGCTGAATTGAAAAGTAAA
GATGAATATAGAATTTTGGCTAAGGGGCAAGGGTTAACAACCCAGCTTTTAGGGTTAAAGACAAGTTCAGTGTCTTCATTT
GAGAAAATGGTTGAGGAAACAAGAGAAAGTATTAAGTCTCAAGAAAGACAAACCATTAAGATAAAATAA
>Components AcGFP1 P4M

**pCMV-GalT-mCherry**
>Amino acid sequence
MRLREPLLSGSAAMPGASLQRACRLLVAVCALHLGVTLVYYLAGRDLSRLPQLVGVSTPLQGGSNSAAAIGQSSGELRTGG
AKDPPVMVSKGEEDNMAIIKEFMRFKVHMEGSVNGHEFEIEGEGEGRPYEGTQTAKLKVTKGGPLPFAWDILSPQFMYGSK

```
AYVKHPADIPDYLKLSFPEGFKWERVMNFEDGGVVTVTQDSSLQDGEFIYKVKLRGTNFPSDGPVMQKKTMGWEASSERMY
PEDGALKGEIKQRLKLKDGGHYDAEVKTTYKAKKPVQLPGAYNVNIKLDITSHNEDYTIVEQYERAEGRHSTGGMDELYK*
>DNA sequence
ATGAGGCTTCGGGAGCCGCTCCTGAGCGGCAGCGCCGCGATGCCAGGCGCGTCCCTACAGCGGGCCTGCCGCCTGCTCGTG
GCCGTCTGCGCTCTGCACCTTGGCGTCACCCTCGTTTACTACCTGGCTGGCCGCGACCTGAGCCGCCTGCCCCAACTGGTC
GGAGTCTCCACACCGCTGCAGGGCGGCTCGAACAGTGCCGCCGCCATCGGGCAGTCCTCCGGGGAGCTCCGGACCGGAGGG
GCCAAGGATCCACCGGTCATGGTGAGCAAGGGCGAGGAGGATAACATGGCCATCATCAAGGAGTTCATGCGCTTCAAGGTG
CACATGGAGGGCTCCGTGAACGGCCACGAGTTCGAGATCGAGGGCGAGGGCGAGGGCCGCCCCTACGAGGGCACCCAGACC
GCCAAGCTGAAGGTGACCAAGGGTGGCCCCCTGCCCTTCGCCTGGGACATCCTGTCCCCTCAGTTCATGTACGGCTCCAAG
GCCTACGTGAAGCACCCCGCCGACATCCCCGACTACTTGAAGCTGTCCTTCCCCGAGGGCTTCAAGTGGGAGCGCGTGATG
AACTTCGAGGACGGCGGCGTGGTGACCGTGACCCAGGACTCCTCCCTGCAGGACGGCGAGTTCATCTACAAGGTGAAGCTG
CGCGGCACCAACTTCCCCTCCGACGGCCCCGTAATGCAGAAGAAGACCATGGGCTGGGAGGCCTCCTCCGAGCGGATGTAC
CCCGAGGACGGCGCCCTGAAGGGCGAGATCAAGCAGAGGCTGAAGCTGAAGGACGGCGGCCACTACGACGCTGAGGTCAAG
ACCACCTACAAGGCCAAGAAGCCCGTGCAGCTGCCCGGCGCCTACAACGTCAACATCAAGTTGGACATCACCTCCCACAAC
GAGGACTACACCATCGTGGAACAGTACGAACGCGCCGAGGGCCGCCACTCCACCGGCGGCATGGACGAGCTGTACAAGTGA
>Components
GalT (N-terminal 81 amino acids of human beta 1,4-galactosyltransferase) mCherry
```

**pCMV-mTagBFP2-FRB-Rab5**

```
>Amino acid sequence
MVSKGEELIKENMHMKLYMEGTVDNHHFKCTSEGEGKPYEGTQTMRIKVVEGGPLPFAFDILATSFLYGSKTFINHTQGIP
DFFKQSFPEGFTWERVTTYEDGGVLTATQDTSLQDGCLIYNVKIRGVNFTSNGPVMQKKTLGWEAFTETLYPADGGLEGRN
DMALKLVGGSHLIANAKTTYRSKKPAKNLKMPGVYYVDYRLERIKEANNETYVEQHEVAVARYCDLPSKLGHKLNSGLRSG
GGGSGGGGSGGGGSRAGGAGAILSRILWHEMWHEGLEEASRLYFGERNVKGMFEVLEPLHAMMERGPQTLKETSFNQAYGR
DLMEAQEWCRKYMKSGNVKDLTQAWDLYYHVFRRISKGGSAGGSAQASNSAVDGTMANRGATRPNGPNTGNKICQFKLVLL
GESAVGKSSLVLRFVKGQFHEFQESTIGAAFLTQTVCLDDTTVKFEIWDTAGQERYHSLAPMYYRGAQAAIVVYDITNEES
FARAKNWVKELQRQASPNIVIALSGNKADLANKRAVDFQEAQSYADDNSLLFMETSAKTSMNVNEIFMAIAKKLPKNEPQN
PGANSARGRGVDLTEPTQPTRSQCCSN*
>DNA sequence
ATGGTGTCTAAGGGCGAAGAGCTGATTAAGGAGAACATGCACATGAAGCTGTACATGGAGGGCACCGTGGACAACCATCAC
TTCAAGTGCACATCCGAGGGCGAAGGCAAGCCCTACGAGGGCACCCAGACCATGAGAATCAAGGTGGTCGAGGGCGGCCCT
CTCCCCTTCGCCTTCGACATCCTGGCTACTAGCTTCCTCTACGGCAGCAAGACCTTCATCAACCACACCCAGGGCATCCCC
GACTTCTTCAAGCAGTCCTTCCCTGAGGGCTTCACATGGGAGAGAGTCACCACATACGAAGACGGGGGCGTGCTGACCGCT
ACCCAGGACACCAGCCTCCAGGACGGCTGCCTCATCTACAACGTCAAGATCAGAGGGGTGAACTTCACATCCAACGGCCCT
GTGATGCAGAAGAAAACACTCGGCTGGGAGGCCTTCACCGAGACGCTGTACCCCGCTGACGGCGGCCTGGAAGGCAGAAAC
GACATGGCCCTGAAGCTCGTGGGCGGGAGCCATCTGATCGCAAACGCCAAGACCACATATAGATCCAAGAAACCCGCTAAG
AACCTCAAGATGCCTGGCGTCTACTATGTGGACTACAGACTGGAAAGAATCAAGGAGGCCAACAACGAGACCTACGTCGAG
CAGCACGAGGTGGCAGTGGCCAGATACTGCGACCTCCCTAGCAAACTGGGGCACAAGCTTAATTCCGGACTCAGATCTGGT
GGCGGAGGCTCGGGCGGAGGTGGGTCGGGTGGCGGCGGATCTCGAGCTGGAGGTGCTGGTGCTATCCTATCTAGAATCCTC
TGGCATGAGATGTGGCATGAAGGCCTGGAAGAGGCATCTCGTTTGTACTTTGGGGAAAGGAACGTGAAAGGCATGTTTGAG
GTGCTGGAGCCCTTGCATGCTATGATGGAACGGGGCCCCCAGACTCTGAAGGAAACATCCTTTAATCAGGCCTATGGTCGA
GATTTAATGGAGGCCCAAGAGTGGTGCAGGAAGTACATGAAATCAGGGAATGTCAAGGACCTCACCCAAGCCTGGGACCTC
TATTATCATGTGTTCCGACGAATCTCAAAGGGTGGTAGTGCTGGTGGTAGTGCTCAAGCTTCGAATTCTGCAGTCGACGGT
ACCATGGCTAATCGAGGAGCAACAAGACCCAACGGGCCAAATACTGGAAATAAAATATGCCAGTTCAAACTAGTACTTCTG
GGAGAGTCTGCTGTTGGCAAATCAAGCCTAGTGCTTCGTTTTGTGAAGGGCCAATTTCATGAATTTCAAGAGAGTACCATA
GGGGCTGCTTTTCTAACCCAAACTGTGTGTCTTGATGATACAACAGTAAAGTTTGAAATATGGGATACAGCTGGTCAAGAA
CGATACCATAGCTTAGCACCAATGTACTACAGAGGAGCACAAGCAGCCATAGTTGTATATGATATCACAAATGAGGAGTCC
TTTGCCAGAGCCAAAAACTGGGTTAAAGAACTTCAGAGGCAAGCCAGTCCTAACATTGTAATAGCTTTATCAGGAAACAAG
GCTGATCTTGCAAATAAAAGAGCTGTCGATTTCCAGGAAGCACAGTCCTATGCAGATGACAACAGTTTATTATTCATGGAG
ACATCAGCTAAAACATCGATGAACGTAAATGAAATATTCATGGCAATAGCTAAAAAGTTGCCAAAGAACGAACCACAGAAT
CCAGGAGCAAATTCTGCCAGAGGAAGAGGAGTAGACCTTACTGAACCCACGCAGCCAACCAGGAGTCAGTGTTGTAGTAAC
TAA
>Components mTagBFP2 FRB Rab5
```

**pCMV-mTagBFP2-FRB-Rab7**

```
>Amino acid sequence
MVSKGEELIKENMHMKLYMEGTVDNHHFKCTSEGEGKPYEGTQTMRIKVVEGGPLPFAFDILATSFLYGSKTFINHTQGIP
DFFKQSFPEGFTWERVTTYEDGGVLTATQDTSLQDGCLIYNVKIRGVNFTSNGPVMQKKTLGWEAFTETLYPADGGLEGRN
DMALKLVGGSHLIANAKTTYRSKKPAKNLKMPGVYYVDYRLERIKEANNETYVEQHEVAVARYCDLPSKLGHKLNSGLRSG
GGGSGGGGSGGGGSRAGGAGAILSRILWHEMWHEGLEEASRLYFGERNVKGMFEVLEPLHAMMERGPQTLKETSFNQAYGR
DLMEAQEWCRKYMKSGNVKDLTQAWDLYYHVFRRISKGGSAGGSAQASMTSRKKVLLKVIILGDSGVGKTSLMNQYVNKKF
SNQYKATIGADFLTKEVMVDDRLVTMQIWDTAGQERFQSLGVAFYRGADCCVLVFDVTAPNTFKTLDSWRDEFLIQASPRD
PENFPFVVLGNKIDLENRQVATKRAQAWCYSKNNIPYFETSAKEAINVEQAFQTIARNALKQETEVELYNEFPEPIKLDKN
DRAKTSAESCSC*
>DNA sequence
ATGGTGTCTAAGGGCGAAGAGCTGATTAAGGAGAACATGCACATGAAGCTGTACATGGAGGGCACCGTGGACAACCATCAC
TTCAAGTGCACATCCGAGGGCGAAGGCAAGCCCTACGAGGGCACCCAGACCATGAGAATCAAGGTGGTCGAGGGCGGCCCT
```

CTCCCCTTCGCCTTCGACATCCTGGCTACTAGCTTCCTCTACGGCAGCAAGACCTTCATCAACCACACCCAGGGCATCCCC
GACTTCTTCAAGCAGTCCTTCCCTGAGGGCTTCACATGGGAGAGAGTCACCACATACGAAGACGGGGGCGTGCTGACCGCT
ACCCAGGACACCAGCCTCCAGGACGGCTGCCTCATCTACAACGTCAAGATCAGAGGGGTGAACTTCACATCCAACGGCCCT
GTGATGCAGAAGAAAACACTCGGCTGGGAGGCCTTCACCGAGACGCTGTACCCCGCTGACGGCGGCCTGGAAGGCAGAAAC
GACATGGCCCTGAAGCTCGTGGGCGGGAGCCATCTGATCGCAAACGCCAAGACCACATATAGATCCAAGAAACCCGCTAAG
AACCTCAAGATGCCTGGCGTCTACTATGTGGACTACAGACTGGAAAGAATCAAGGAGGCCAACAACGAGACCTACGTCGAG
CAGCACGAGGTGGCAGTGGCCAGATACTGCGACCTCCCTAGCAAACTGGGGCACAAGCTTAATTCCGGACTCAGATCTGGT
GGCGGAGGCTCGGGCGGAGGTGGGTCGGGTGGCGGCGGATCTCGAGCTGGAGGTGCTGGTGCTATCCTATCTAGAATCCTC
TGGCATGAGATGTGGCATGAAGGCCTGGAAGAGGCATCTCGTTTGTACTTTGGGGAAAGGAACGTGAAAGGCATGTTTGAG
GTGCTGGAGCCCTTGCATGCTATGATGGAACGGGGCCCCCAGACTCTGAAGGAAACATCCTTTAATCAGGCCTATGGTCGA
GATTTAATGGAGGCCCAAGAGTGGTGCAGGAAGTACATGAAATCAGGGAATGTCAAGGACCTCACCCAAGCCTGGGACCTC
TATTATCATGTGTTCCGACGAATCTCAAAGGGTGGTAGTGCTGGTGGTAGTGCTCAAGCTTCGATGACCTCTAGGAAGAAA
GTGTTGCTGAAGGTTATCATCCTGGGAGATTCTGGAGTTGGTAAGACATCACTCATGAACCAGTATGTGAACAAGAAATTC
AGTAATCAGTACAAAGCTACAATAGGAGCAGACTTTCTGACAAAGGAGGTGATGGTGGATGACAGACTAGTTACAATGCAG
ATCTGGGACACAGCAGGCCAGGAACGGTTCCAGTCCCTTGGTGTGGCCTTCTACAGAGGTGCAGACTGCTGCGTTCTGGTA
TTTGACGTTACTGCCCCCAACACATTCAAAACCCTCGATAGCTGGAGAGATGAGTTTCTCATCCAGGCCAGTCCCCGGGAT
CCTGAAAACTTCCCTTTCGTTGTGTTGGGAAACAAGATTGACCTCGAAAACAGACAAGTGGCCACAAAGCGGGCACAGGCC
TGGTGCTACAGCAAAAACAACATTCCCTACTTCGAGACCAGTGCCAAGGAGGCCATCAATGTGGAGCAGGCGTTCCAGACG
ATTGCAAGGAATGCACTTAAACAGGAAACAGAGGTGGAGCTGTACAATGAATTCCCTGAACCCATCAAACTGGACAAGAAC
GACCGGGCCAAGACCTCAGCGGAAAGCTGCAGTTGCTGA
>Components mTagBFP2 FRB Rab7

**pCMV-LAMP1-mCherry**
>Amino acid sequence
MAAPGARRPLLLLLLAGLAHSAPALFEVKDNNGTACIMASFSASFLTTYEAGHVSKVSNMTLPASAEVLKNSSSCGEKNAS
EPTLAITFGEGYLLKLTFTKNTTRYSVQHMYFTYNLSDTQFFPNASSKGPDTVDSTTDIKADINKTYRCVSDIRVYMKNVT
IVLWDATIQAYLPSSNFSKEETRCPQDQPSPTTGPPSPSPPLVPTNPSVSKYNVTGDNGTCLLASMALQLNITYMKKDNTT
VTRAFNINPSDKYSGTCGAQLVTLKVGNKSRVLELQFGMNATSSLFFLQGVQLNMTLPDAIEPTFSTSNYSLKALQASVGN
SYKCNSEEHIFVSKALALNVFSVQVQAFRVESDRFGSVEECVQDGNNMLIPIAVGGALAGLVLIVLIAYLIGRKRSHAGYQ
TISEFGSTGSTGSTGADPPVATMVSKGEEDNMAIIKEFMRFKVHMEGSVNGHEFEIEGEGEGRPYEGTQTAKLKVTKGGPL
PFAWDILSPQFMYGSKAYVKHPADIPDYLKLSFPEGFKWERVMNFEDGGVVTVTQDSSLQDGEFIYKVKLRGTNFPSDGPV
MQKKTMGWEASSERMYPEDGALKGEIKQRLKLKDGGHYDAEVKTTYKAKKPVQLPGAYNVNIKLDITSHNEDYTIVEQYER
AEGRHSTGGMDELYK*
>DNA sequence
ATGGCGGCCCCGGGCGCCCGGCGGCCGCTGCTCCTGTTGCTGCTGGCAGGCCTTGCACACAGCGCCCCAGCACTGTTCGAG
GTGAAAGACAACAACGGCACAGCGTGTATAATGGCCAGCTTCTCTGCCTCCTTTCTGACCACCTATGAGGCTGGACATGTT
TCTAAGGTCTCGAATATGACCCTGCCAGCCTCTGCAGAAGTCCTGAAGAATAGCAGCTCTTGTGGTGAAAAGAATGCTTCT
GAGCCCACCCTCGCAATCACCTTTGGAGAAGGATATTTACTGAAACTCACCTTCACAAAAAACACAACACGTTACAGTGTC
CAGCACATGTATTTCACATATAACCTGTCAGACACACAATTCTTTCCCAATGCCAGCTCCAAAGGGCCCGACACTGTGGAT
TCCACAACTGACATCAAGGCAGACATCAACAAAACATACCGATGTGTCAGCGACATCAGGGTCTACATGAAGAATGTGACC
ATTGTGCTCTGGGACGCTACTATCCAGGCCTACCTGCCGAGTAGCAACTTCAGCAAGGAAGAGACACGCTGCCCACAGGAT
CAACCTTCCCCAACTACTGGGCCACCCAGCCCCTCACCACCACTTGTGCCCACAAACCCCAGTGTGTCCAAGTACAATGTG
ACTGGTGACAATGGAACCTGCCTGCTGGCCTCTATGGCACTGCAACTCAACATCACCTACATGAAGAAGGACAACACGACT
GTGACCAGAGCATTCAACATCAACCCAAGTGACAAATATAGTGGGACTTGCGGTGCCCAGTTGGTGACCCTGAAGGTGGGG
AACAAGAGCAGAGTCCTGGAGCTGCAGTTTGGGATGAATGCCACTTCTAGCCTGTTTTTCCTGCAAGGAGTTCAGTTGAAC
ATGACTCTTCCTGATGCCATAGAGCCCACGTTCAGCACCTCCAACTATTCCCTGAAAGCTCTTCAGGCCAGTGTCGGCAAC
TCATACAAGTGCAACAGTGAGGAGCACATCTTTGTCAGCAAGGCGCTCGCCCTCAATGTCTTCAGCGTGCAAGTCCAGGCT
TTCAGGGTAGAAAGTGACAGGTTTGGGTCTGTGGAAGAGTGTGTACAGGACGGTAACAACATGCTGATCCCCATTGCTGTG
GGCGGGGCCCTGGCAGGGCTGGTCCTCATCGTCCTCATCGCCTACCTCATCGGCAGGAAGAGGAGTCACGCGGGCTATCAG
ACCATCTCGGAATTCGGCTCCACCGGCTCCACCGGCTCCACCGGCGCGGATCCACCGGTCGCCACCATGGTGAGCAAGGGC
GAGGAGGATAACATGGCCATCATCAAGGAGTTCATGCGCTTCAAGGTGCACATGGAGGGCTCCGTGAACGGCCACGAGTTC
GAGATCGAGGGCGAGGGCGAGGGCCGCCCCTACGAGGGCACCCAGACCGCCAAGCTGAAGGTGACCAAGGGTGGCCCCCTG
CCCTTCGCCTGGGACATCCTGTCCCCTCAGTTCATGTACGGCTCCAAGGCCTACGTGAAGCACCCCGCCGACATCCCCGAC
TACTTGAAGCTGTCCTTCCCCGAGGGCTTCAAGTGGGAGCGCGTGATGAACTTCGAGGACGGCGGCGTGGTGACCGTGACC
CAGGACTCCTCCCTGCAGGACGGCGAGTTCATCTACAAGGTGAAGCTGCGCGGCACCAACTTCCCCTCCGACGGCCCCGTA
ATGCAGAAGAAGACCATGGGCTGGGAGGCCTCCTCCGAGCGGATGTACCCCGAGGACGGCGCCCTGAAGGGCGAGATCAAG
CAGAGGCTGAAGCTGAAGGACGGCGGCCACTACGACGCTGAGGTCAAGACCACCTACAAGGCCAAGAAGCCCGTGCAGCTG
CCCGGCGCCTACAACGTCAACATCAAGTTGGACATCACCTCCCACAACGAGGACTACACCATCGTGGAACAGTACGAACGC
GCCGAGGGCCGCCACTCCACCGGCGGCATGGACGAGCTGTACAAGTAA
>Components LAMP1 mCherry

**pCMV-mTagBFP2-PLCδ-PH**
>Amino acid sequence
MVSKGEELIKENMHMKLYMEGTVDNHHFKCTSEGEGKPYEGTQTMRIKVVEGGPLPFAFDILATSFLYGSKTFINHTQGIP
DFFKQSFPEGFTWERVTTYEDGGVLTATQDTSLQDGCLIYNVKIRGVNFTSNGPVMQKKTLGWEAFTETLYPADGGLEGRN
DMALKLVGGSHLIANAKTTYRSKKPAKNLKMPGVYYVDYRLERIKEANNETYVEQHEVAVARYCDLPSKLGHKLNSGLRSR
AQASNSAVDGTAGPGSMDSGRDFLTLHGLQDDEDLQALLKGSQLLKVKSSSWRRERFYKLQEDCKTIWQESRKVMRTPESQ

LFSIEDIQEVRMGHRTEGLEKFARDVPEDRCFSIVFKDQRNTLDLIAPSPADAQHWVLGLHKIIHHSGSMDQRQKLQHWIH
SCLRKADKNKDNKMSFKELQNFLKELNIQ*
>DNA sequence
ATGGTGTCTAAGGGCGAAGAGCTGATTAAGGAGAACATGCACATGAAGCTGTACATGGAGGGCACCGTGGACAACCATCAC
TTCAAGTGCACATCCGAGGGCGAAGGCAAGCCCTACGAGGGCACCCAGACCATGAGAATCAAGGTGGTCGAGGGCGGCCCT
CTCCCCTTCGCCTTCGACATCCTGGCTACTAGCTTCCTCTACGGCAGCAAGACCTTCATCAACCACACCCAGGGCATCCCC
GACTTCTTCAAGCAGTCCTTCCCTGAGGGCTTCACATGGGAGAGAGTCACCACATACGAAGACGGGGGCGTGCTGACCGCT
ACCCAGGACACCAGCCTCCAGGACGGCTGCCTCATCTACAACGTCAAGATCAGAGGGGTGAACTTCACATCCAACGGCCCT
GTGATGCAGAAGAAAACACTCGGCTGGGAGGCCTTCACCGAGACGCTGTACCCCGCTGACGGCGGCCTGGAAGGCAGAAAC
GACATGGCCCTGAAGCTCGTGGGCGGGAGCCATCTGATCGCAAACGCCAAGACCACATATAGATCCAAGAAACCCGCTAAG
AACCTCAAGATGCCTGGCGTCTACTATGTGGACTACAGACTGGAAAGAATCAAGGAGGCCAACAACGAGACCTACGTCGAG
CAGCACGAGGTGGCAGTGGCCAGATACTGCGACCTCCCTAGCAAACTGGGGCACAAGCTTAATTCCGGACTCAGATCTCGA
GCTCAAGCTTCGAATTCTGCAGTCGACGGTACCGCGGGCCCGGGATCCATGGACTCGGGCCGGGACTTCCTGACCCTGCAC
GGCCTACAGGATGATGAGGATCTACAGGCGCTGCTGAAGGGCAGCCAGCTCCTGAAGGTGAAGTCCAGCTCATGGAGGAGA
GAGCGGTTCTACAAGTTGCAGGAGGACTGCAAGACCATCTGGCAGGAGTCCCGCAAGGTCATGCGGACCCCGGAGTCCCAG
CTGTTCTCCATCGAGGACATTCAGGAGGTGCGAATGGGGCACCGCACGGAGGGTCTGGAGAAGTTCGCCCGTGATGTGCCC
GAGGACCGCTGCTTCTCCATTGTCTTCAAGGACCAGCGCAATACACTAGACCTCATCGCCCCATCGCCAGCTGATGCCCAG
CACTGGGTGCTGGGGCTGCACAAGATCATCCACCACTCAGGCTCCATGGACCAGCGTCAGAAGCTACAGCACTGGATTCAC
TCCTGCTTGCGAAAAGCTGACAAAAACAAGGACAACAAGATGAGCTTCAAGGAGCTGCAGAACTTCCTGAAGGAGCTCAAC
ATCCAGTAA
>Components mTagBFP2 PLCδ-PH

**pCAGGS-HaloTag-KRas4B(CT)**
>Amino acid sequence
MAEIGTGFPFDPHYVEVLGERMHYVDVGPRDGTPVLFLHGNPTSSYVWRNIIPHVAPTHRCIAPDLIGMGKSDKPDLGYFF
DDHVRFMDAFIEALGLEEVVLVIHDWGSALGFHWAKRNPERVKGIAFMEFIRPIPTWDEWPEFARETFQAFRTTDVGRKLI
IDQNVFIEGTLPMGVVRPLTEVEMDHYREPFLNPVDREPLWRFPNELPIAGEPANIVALVEEYMDWLHQSPVPKLLFWGTP
GVLIPPAEAARLAKSLPNCKAVDIGPGLNLLQEDNPDLIGSEIARWLSTLEISGGSGASAGGGSLEISGGKKKKKKSKTKC
VIM*
>DNA sequence
ATGGCAGAAATCGGTACTGGCTTTCCATTCGACCCCCATTATGTGGAAGTCCTGGGCGAGCGCATGCACTACGTCGATGTT
GGTCCGCGCGATGGCACCCCTGTGCTGTTCCTGCACGGTAACCCGACCTCCTCCTACGTGTGGCGCAACATCATCCCGCAT
GTTGCACCGACCCATCGCTGCATTGCTCCAGACCTGATCGGTATGGGCAAATCCGACAAACCAGACCTGGGTTATTTCTTC
GACGACCACGTCCGCTTCATGGATGCCTTCATCGAAGCCCTGGGTCTGGAAGAGGTCGTCCTGGTCATTCACGACTGGGGC
TCCGCTCTGGGTTTCCACTGGGCCAAGCGCAATCCAGAGCGCGTCAAAGGTATTGCATTTATGGAGTTCATCCGCCCTATC
CCGACCTGGGACGAATGGCCAGAATTTGCCCGCGAGACCTTCCAGGCCTTCCGCACCACCGACGTCGGCCGCAAGCTGATC
ATCGATCAGAACGTTTTTATCGAGGGTACGCTGCCGATGGGTGTCGTCCGCCCGCTGACTGAAGTCGAGATGGACCATTAC
CGCGAGCCGTTCCTGAATCCTGTTGACCGCGAGCCACTGTGGCGCTTCCCAAACGAGCTGCCAATCGCCGGTGAGCCAGCG
AACATCGTCGCGCTGGTCGAAGAATACATGGACTGGCTGCACCAGTCCCCTGTCCCGAAGCTGCTGTTCTGGGGCACCCCA
GGCGTTCTGATCCCACCGGCCGAAGCCGCTCGCCTGGCCAAAAGCCTGCCTAACTGCAAGGCTGTGGACATCGGCCCGGGT
CTGAATCTGCTGCAAGAAGACAACCCGGACCTGATCGGCAGCGAGATCGCGCGCTGGCTGTCGACGCTCGAGATTTCCGGC
GGCTCCGGTGCCAGTGCTGGTGGTGGCAGCCTCGAGATTTCCGGCGGGAAGAAAAAGAAGAAGAAGTCCAAGACAAAATGC
GTGATTATGTAG
>Components HaloTag KRas4B(CT)

**pCMV-mScarlet-I-eDHFR-ORP5(ΔPH)**
>Amino acid sequence
MVSKGEAVIKEFMRFKVHMEGSMNGHEFEIEGEGEGRPYEGTQTAKLKVTKGGPLPFSWDILSPQFMYGSRAFTKHPADIP
DYYKQSFPEGFKWERVMNFEDGGAVTVTQDTSLEDGTLIYKVKLRGTNFPPDGPVMQKKTMGWEASTERLYPEDGVLKGDI
KMALRLKDGGRYLADFKTTYKAKKPVQMPGAYNVDRKLDITSHNEDYTVVEQYERSEGRHSTGGMDELYKSGLRSRSAAAG
AGGAARAAMISLIAALAVDRVIGMENAMPWNLPADLAWFKRNTLNKPVIMGRHTWESIGRPLPGRKNIILSSQPGTDDRVT
WVKSVDEAIAACGDVPEIMVIGGGRVYEQFLPKAQKLYLTHIDAEVEGDTHFPDYEPDDWESVFSEFHDADAQNSHSYCFE
ILERRAGSGGGTGAGGSGGGSRAQASCKPGRDGEPGTSPDASPSSLCGLPASATVHPDQDLFPLNGSSLENDAFSDKSERE
NPEESDTETQDHSRKTESGSDQSETPGAPVRRGTTYVEQVQEELGELGEASQVETVSEENKSLMWTLLKQLRPGMDLSRVV
LPTFVLEPRSFLNKLSDYYYHADLLSRAAVEEDAYSRMKLVLRWYLSGFYKKPKGIKKPYNPILGETFRCCWFHPQTDSRT
FYIAEQVSHHPPVSAFHVSNRKDGFCISGSITAKSRFYGNSLSALLDGKATLTFLNRAEDYTLTMPYAHCKGILYGTMTLE
LGGKVTIECAKNNFQAQLEFKLKPFFGGSTSINQISGKITSGEEVLASLSGHWDRDVFIKEEGSGSSALFWTPSGEVRRQR
LRQHTVPLEEQTELESERLWQHVTRAISKGDQHRATQEKFALEEAQRQRARERQESLMPWKPQLFHLDPITQEWHYRYEDH
SPWDPLKDIAQFEQDGILRTLQQEAVARQTTFLGSPGPRHERSGPDQRLRKASDQPSGHSQATESSGSTPESCPELSDEEQ
DGDFVPGGESPCPRCRKEARRLQALHEAILSIREAQQELHRHLSAMLSSTARAAQAPTPGLLQSPRSWFLLCVFLACQLFI
NHILK*
>DNA sequence
ATGGTGAGCAAGGGCGAGGCAGTGATCAAGGAGTTCATGCGGTTCAAGGTGCACATGGAGGGCTCCATGAACGGCCACGAG
TTCGAGATCGAGGGCGAGGGCGAGGGCCGCCCCTACGAGGGCACCCAGACCGCCAAGCTGAAGGTGACCAAGGGTGGCCCC
CTGCCCTTCTCCTGGGACATCCTGTCCCCTCAGTTCATGTACGGCTCCAGGGCCTTCACCAAGCACCCCGCCGACATCCCC
GACTACTATAAGCAGTCCTTCCCCGAGGGCTTCAAGTGGGAGCGCGTGATGAACTTCGAGGACGGCGGCGCCGTGACCGTG
ACCCAGGACACCTCCCTGGAGGACGGCACCCTGATCTACAAGGTGAAGCTCCGCGGCACCAACTTCCCTCCTGACGGCCCC

GTAATGCAGAAGAAGACAATGGGCTGGGAAGCGTCCACCGAGCGGTTGTACCCCGAGGACGGCGTGCTGAAGGGCGACATT
AAGATGGCCCTGCGCCTGAAGGACGGCGGCCGCTACCTGGCGGACTTCAAGACCACCTACAAGGCCAAGAAGCCCGTGCAG
ATGCCCGGCGCCTACAACGTCGACCGCAAGTTGGACATCACCTCCCACAACGAGGACTACACCGTGGTGGAACAGTACGAA
CGCTCCGAGGGCCGCCACTCCACCGGCGGCATGGACGAGCTGTACAAGTCCGGACTCAGATCTCGAAGCGCGGCCGCGGGA
GCAGGAGGAGCAGCTCGAGCGGCGATGATTTCTCTGATTGCCGCTCTGGCCGTGGACAGAGTGATCGGCATGGAAAACGCC
ATGCCTTGGAACCTGCCTGCCGATCTGGCCTGGTTCAAGCGGAATACCCTGAACAAGCCCGTGATCATGGGCAGACACACC
TGGGAGTCTATCGGCAGACCTCTGCCTGGCCGGAAGAACATCATTCTGTCTAGCCAGCCTGGCACCGACGATAGAGTGACA
TGGGTCAAGAGCGTGGACGAGGCCATTGCTGCTTGCGGAGATGTGCCTGAGATCATGGTTATCGGCGGAGGCCGGGTGTAC
GAGCAGTTTCTGCCTAAGGCTCAGAAGCTGTACCTGACACACATCGACGCCGAGGTGGAAGGCGATACACACTTCCCCGAC
TACGAGCCCGATGACTGGGAGAGCGTGTTCAGCGAGTTTCACGACGCCGACGCTCAGAACAGCCACAGCTACTGCTTCGAG
ATCCTGGAAAGACGCGCCGGATCTGGGGGCGGTACCGGAGCTGGAGGGAGCGGCGGGGGAAGTCGAGCTCAAGCTTCGTGC
AAGCCGGGCCGAGACGGGGAGCCAGGGACCTCGCCAGACGCATCACCCTCATCGCTCTGTGGGCTGCCAGCCTCAGCCACC
GTCCACCCAGACCAAGACCTGTTCCCACTGAACGGGTCTTCCCTGGAGAACGATGCATTCTCAGACAAGTCGGAGAGAGAG
AACCCTGAGGAGTCAGATACCGAGACCCAGGACCATAGCCGGAAGACGGAGAGTGGCAGCGACCAGTCAGAGACCCCTGGG
GCCCCGGTGCGGAGAGGGACCACCTATGTGGAGCAGGTCCAGGAGGAGCTGGGGGAGCTGGGCGAGGCGTCCCAGGTGGAG
ACAGTGTCAGAGGAGAACAAGAGTCTGATGTGGACCCTGCTGAAGCAGCTACGGCCAGGCATGGACCTGTCCCGCGTGGTG
CTACCCACGTTCGTACTGGAGCCGCGCTCCTTCCTGAACAAGCTCTCCGACTACTACTACCACGCAGACCTGCTCTCCAGG
GCTGCGGTGGAGGAGGATGCCTACAGCCGCATGAAGCTGGTGCTGCGGTGGTACCTGTCTGGCTTCTACAAGAAGCCCAAG
GGAATCAAGAAGCCGTACAACCCCATCCTGGGGGAGACCTTCCGCTGCTGCTGGTTCCACCCGCAGACTGACAGCCGCACA
TTCTACATAGCAGAGCAGGTGTCCCACCACCCGCCCGTGTCTGCCTTCCACGTCAGCAACCGGAAGGACGGCTTCTGCATC
AGTGGCAGCATCACAGCCAAGTCCAGGTTTTATGGAACTCGCTGTCGGCGCTGCTGGACGGCAAAGCCACGCTCACCTTC
CTGAACCGAGCCGAGGATTACACCCTTACCATGCCCTACGCCCACTGCAAAGGAATCCTGTATGGCACGATGACCCTGGAG
CTGGGTGGAAGGTCACCATCGAGTGTGCGAAGAACAACTTCCAGGCCCAGCTGGAATTCAAACTCAAGCCCTTCTTCGGG
GGTAGCACCAGCATCAACCAGATCTCGGGAAAGATCACGTCGGGAGAGGAAGTCCTGGCGAGCCTCAGTGGCCACTGGGAC
AGGGACGTGTTTATCAAGGAGGAAGGGAGCGGAAGCAGTGCGCTTTTCTGGACCCCGAGCGGGGAGGTCCGCAGACAGAGG
CTGAGGCAGCACACGGTGCCGCTGGAGGAGCAGACGGAGCTGGAGTCCGAGAGGCTCTGGCAGCACGTCACCAGGGCCATC
AGCAAGGGCGACCAGCACAGGGCCACACAGGAGAAGTTTGCACTGGAGGAGGCACAGCGGCAGCGGGCCCGTGAGCGGCAG
GAGAGCCTCATGCCCTGGAAGCCGCAGCTGTTCCACCTGGACCCCATCACCCAGGAGTGGCACTACCGATACGAGGACCAC
AGCCCCTGGGACCCCCTGAAGGACATCGCCCAGTTTGAGCAAGACGGGATCCTGCGGACCTTGCAGCAGGAGGCCGTGGCC
CGCCAGACCACCTTCCTGGGCAGCCCAGGGCCCAGGCACGAGAGGTCTGGCCCAGACCAGCGGCTTCGCAAGGCCAGCGAC
CAGCCCTCCGGCCACAGCCAGGCCACGGAGAGCAGCGGATCCACGCCTGAGTCCTGCCCAGAGCTCTCAGACGAGGAGCAG
GATGGTGACTTTGTCCCTGGCGGTGAGAGCCCATGCCCTCGGTGCAGGAAGGAGGCGCGGCGGCTGCAGGCCCTGCACGAG
GCCATCCTCTCCATCCGAGAGGCCCAGCAGGAGCTGCACAGGCACCTCTCGGCCATGCTGAGCTCCACGGCACGGGCAGCA
CAGGCACCGACCCCAGGCCTCCTGCAGAGCCCCCGATCCTGGTTCCTGCTCTGCGTGTTCCTGGCGTGTCAGCTGTTCATT
AACCACATCCTCAAATAG
>Components mScarlet-I eDHFR ORP5(ΔPH)

**pT5-ORP9-PH-NusA-His6**

>Amino acid sequence
MTNRSGSASIVEGPLSKWTNVMKGWQYRWFVLDYNAGLLSYYTSKDKMMRGSRRGCVRLRGAVIGIDDEDDSTFTITVDQK
TFHFQARDADEREKWIHALEETILRHTLQLQGLDSGTSGGGGENLYFQGSGELNKEILAVVEAVSNEKALPREKIFEALES
ALATATKKKYEQEIDVRVQIDRKSGDFDTFRRWLVVDEVTQPTKEITLEAARYEDESLNLGDYVEDQIESVTFDRITTQTA
KQVIVQKVREAERAMVVDQFREHEGEIITGVVKKVNRDNISLDLGNNAEAVILREDMLPRENFRPGDRVRGVLYSVRPEAR
GAQLFVTRSKPEMLIELFRIEVPEIGEEVIEIKAAARDPGSRAKIAVKTNDKRIDPVGACVGMRGARVQAVSTELGGERID
IVLWDDNPAQFVINAMAPADVASIVVDEDKHTMDIAVEAGNLAQAIGRNGQNVRLASQLSGWELNVMTVDDLQAKHQAEAH
AAIDTFTKYLDIDEDFATVLVEEGFSTLEELAYVPMKELLEIEGLDEPTVEALRERAKNALATIAQAQEESLGDNKPADDL
LNLEGVDRDLAFKLAARGVCTLEDLAEQGIDDLADIEGLTDEKAGALIMAARNICWFGDEAGSGSGSGSLPETGGGSGHHH
HHH*
>DNA sequence
ATGACAAATCGATCTGGTTCTGCGTCGATCGTTGAAGGCCCGCTGAGCAAATGGACCAACGTAATGAAGGGTTGGCAGTAT
CGTTGGTTCGTGCTGGACTACAATGCGGGGCTGCTCTCTTACTATACGTCCAAAGATAAAATGATGCGCGGCAGTCGCCGT
GGCTGCGTTCGCCTGCGGGGCGCTGTGATTGGTATTGACGACGAAGATGACAGCACCTTTACAATTACCGTCGATCAGAAA
ACCTTTCACTTCCAGGCACGTGATGCAGATGAGCGAGAGAAGTGGATTCATGCCTTAGAAGAAACTATTCTGCGCCATACG
TTACAGCTTCAAGGTTTGGATTCAGGAACTAGTGGCGGCGGAGGAGAGAACCTGTACTTTCAGGGTTCAGGAGAGCTCAAC
AAAGAAATTTTGGCTGTAGTTGAAGCCGTATCCAATGAAAAGGCGCTACCTCGCGAGAAGATTTTCGAAGCATTGGAAAGC
GCGCTGGCGACAGCAACAAAGAAAAAATATGAACAAGAGATCGACGTCCGCGTACAGATCGATCGCAAAAGCGGTGATTTT
GACACTTTCCGTCGCTGGTTAGTTGTTGATGAAGTCACCCAGCCGACCAAGGAAATCACCCTTGAAGCCGCACGTTATGAA
GATGAAAGCCTGAACCTGGGCGATTACGTTGAAGATCAGATTGAGTCTGTTACCTTTGACCGTATCACTACCCAGACGGCA
AAACAGGTTATCGTGCAGAAAGTGCGTGAAGCCGAACGTGCGATGGTGGTTGATCAGTTCCGTGAACACGAAGGTGAAATC
ATCACCGGCGTGGTGAAAAAAGTAAACCGCGACAACATCTCTCTGGATCTGGGCAACAACGCTGAAGCCGTGATCCTGCGC
GAAGATATGCTGCCGCGTGAAAACTTCCGCCCTGGCGACCGCGTTCGTGGCGTGCTCTATTCCGTTCGCCCGGAAGCGCGT
GGCGCGCAACTGTTCGTCACTCGTTCCAAGCCGGAAATGCTGATCGAACTGTTCCGTATTGAAGTGCCAGAAATCGGCGAA
GAAGTGATTGAAATTAAAGCAGCGGCTCGCGATCCGGGTTCTCGTGCGAAAATCGCGGTGAAAACCAACGATAAACGTATC
GATCCGGTAGGTGCTTGCGTAGGTATGCGTGGCGCGCGTGTTCAGGCGGTGTCTACTGAACTGGGTGGCGAGCGTATCGAT
ATCGTCCTGTGGGATGATAACCCGGCGCAGTTCGTGATTAACGCAATGGCACCGGCAGACGTTGCTTCTATCGTGGTGGAT
GAAGATAAACACACCATGGACATCGCCGTTGAAGCCGGTAATCTGGCGCAGGCGATTGGCCGTAACGGTCAGAACGTGCGT
CTGGCTTCGCAACTGAGCGGTTGGGAACTCAACGTGATGACCGTTGACGACCTGCAAGCTAAGCATCAGGCGGAAGCGCAC
GCAGCGATCGACACCTTCACCAAATATCTCGACATCGACGAAGACTTCGCGACTGTTCTGGTAGAAGAAGGCTTCTCGACG

```
CTGGAAGAATTGGCCTATGTGCCGATGAAAGAGCTGTTGGAAATCGAAGGCCTTGATGAGCCGACCGTTGAAGCACTGCGC
GAGCGTGCTAAAAATGCACTGGCCACCATTGCACAGGCCCAGGAAGAAAGCCTCGGTGATAACAAACCGGCTGACGATCTG
CTGAACCTTGAAGGGGTAGATCGTGATTTGGCATTCAAACTGGCCGCCCGTGGCGTTTGTACGCTGGAAGATCTCGCCGAA
CAGGGCATTGATGATCTGGCTGATATCGAAGGGTTGACCGACGAAAAAGCCGGAGCACTGATTATGGCTGCCCGTAATATT
TGCTGGTTCGGTGACGAAGCGGGATCCGGTAGTGGATCTGGATCACTGCCGGAAACCGGCGGTGGCAGCGGTCACCATCAT
CACCACCACTAA
```
>Components ORP9-PH TEV protease cleavage sequence NusA His6
